# Identifying Nootropic Drug Targets via Large-Scale Cognitive GWAS and Transcriptomics

**DOI:** 10.1101/2020.02.06.934752

**Authors:** Max Lam, Chen Chia-Yen, Xia Yan, W. David Hill, Joey W. Trampush, Jin Yu, Emma Knowles, Gail Davies, Eli Stahl, Laura Huckins, David C. Liewald, Srdjan Djurovic, Ingrid Melle, Andrea Christoforou, Ivar Reinvang, Pamela DeRosse, Astri J. Lundervold, Vidar M. Steen, Thomas Espeseth, Katri Räikkönen, Elisabeth Widen, Aarno Palotie, Johan G. Eriksson, Ina Giegling, Bettina Konte, Annette M. Hartmann, Panos Roussos, Stella Giakoumaki, Katherine E. Burdick, Antony Payton, William Ollier, Ornit Chiba-Falek, Deborah K. Koltai, Anna C. Need, Elizabeth T. Cirulli, Aristotle N. Voineskos, Nikos C. Stefanis, Dimitrios Avramopoulos, Alex Hatzimanolis, Nikolaos Smyrnis, Robert M. Bilder, Nelson A. Freimer, Tyrone D. Cannon, Edythe London, Russell A. Poldrack, Fred W. Sabb, Eliza Congdon, Emily Drabant Conley, Matthew A. Scult, Dwight Dickinson, Richard E. Straub, Gary Donohoe, Derek Morris, Aiden Corvin, Michael Gill, Ahmad R. Hariri, Daniel R. Weinberger, Neil Pendleton, Panos Bitsios, Dan Rujescu, Jari Lahti, Stephanie Le Hellard, Matthew C. Keller, Ole A. Andreassen, Ian J. Deary, David C. Glahn, Liu Chunyu, Anil K. Malhotra, Todd Lencz

## Abstract

**Background:** Cognitive traits demonstrate significant genetic correlations with many psychiatric disorders and other health-related traits. Many neuropsychiatric and neurodegenerative disorders are marked by cognitive deficits. Therefore, genome-wide association studies (GWAS) of general cognitive ability might suggest potential targets for nootropic drug repurposing. Our previous effort to identify “druggable genes” (i.e., GWAS-identified genes that produce proteins targeted by known small molecules) was modestly powered due to the small cognitive GWAS sample available at the time. Since then, two large cognitive GWAS meta-analyses have reported 148 and 205 genome-wide significant loci, respectively. Additionally, large-scale gene expression databases, derived from post-mortem human brain, have recently been made available for GWAS annotation. Here, we 1) reconcile results from these two cognitive GWAS meta-analyses to further enhance power for locus discovery; 2) employ several complementary transcriptomic methods to identify genes in these loci with variants that are credibly associated with cognition; and 3) further annotate the resulting genes to identify “druggable” targets.

**Methods:** GWAS summary statistics were harmonized and jointly analysed using Multi-Trait Analysis of GWAS [MTAG], which is optimized for handling sample overlaps. Downstream gene identification was carried out using MAGMA, S-PrediXcan/S-TissueXcan Transcriptomic Wide Analysis, and eQTL mapping, as well as more recently developed methods that integrate GWAS and eQTL data via Summary-statistics Mendelian Randomization [SMR] and linkage methods [HEIDI], Available brain-specific eQTL databases included GTEXv7, BrainEAC, CommonMind, ROSMAP, and PsychENCODE. Intersecting credible genes were then annotated against multiple chemoinformatic databases [DGIdb, K_I_, and a published review on “druggability”].

**Results:** Using our meta-analytic data set (N = 373,617) we identified 241 independent cognition-associated loci (29 novel), and 76 genes were identified by 2 or more methods of gene identification. 26 genes were associated with general cognitive ability via SMR, 16 genes via STissueXcan/S-PrediXcan, 47 genes via eQTL mapping, and 68 genes via MAGMA pathway analysis. The use of the HEIDI test permitted the exclusion of candidate genes that may have been artifactually associated to cognition due to linkage, rather than direct causal or indirect pleiotropic effects. Actin and chromatin binding gene sets were identified as novel pathways that could be targeted via drug repurposing. Leveraging on our various transcriptome and pathway analyses, as well as available chemoinformatic databases, we identified 16 putative genes that may suggest drug targets with nootropic properties.

**Discussion:** Results converged on several categories of significant drug targets, including serotonergic and glutamatergic genes, voltage-gated ion channel genes, carbonic anhydrase genes, and phosphodiesterase genes. The current results represent the first efforts to apply a multi-method approach to integrate gene expression and SNP level data to identify credible actionable genes for general cognitive ability.

## Introduction

One central goal for genome-wide association studies (GWAS) is the identification of potential targets for clinically useful pharmacologic interventions; drugs whose targets have supporting genetic evidence of association to the indication are significantly more likely to successfully reach approval than those without such evidence^1–3^. While novel drug targets for major psychiatric illnesses have emerged from recent large-scale GWAS^4–7^, broad-based cognitive deficits are an enduring and disabling feature for many patients with severe mental illness that are inadequately addressed by current medications.^8^. Similarly, effective cognitive enhancing medications (“nootropics”) are limited for patients with dementias and other neurodegenerative disorders^9^. Thus, the genetic study of general cognitive ability (GCA) holds the potential for identifying novel targets for nootropic medications, that could have widespread applications^10^. The genetic architecture of general cognitive ability (GCA) has been examined with increasingly large sample sizes over the last few years^11–17^. Physical health, illness, mortality^18^, and psychiatric traits^19^ have shown significant genetic correlations with individual differences in GCA. Dissecting the pleiotropic genetic architectures underlying GCA, educational attainment, and schizophrenia. We have recently shown that neurodevelopmental pathways and adulthood synaptic processes are dissociable etiologic mechanisms relating to genetic liability to psychosis^20^.

Nevertheless, identifying specific genes functionally linked to GCA, with protein products that could be targeted by pharmacological agents, remains a core challenge. Using a pathway-based methodology^21,22^, we previously reported that several genes encoding T-and L-type calcium channels, targeted by known pharmaceuticals, were associated with GCA^10^; however, that study was relatively underpowered. Now with much larger GWAS of cognition available^15,16^, we can more effectively and reliably identify putative drug targets for further investigation. Contemporaneously, the latest large-scale brain eQTL databases substantially enhance the assignment of regional GWAS signals to specific genes that can then be interrogated for druggability^23–29^. Further, recent advances in genetic epidemiology methods (e.g. Mendelian randomization) have enabled identification of potentially spurious eQTL associations that may be based on linkage rather than meaningful biology.^30^. Thus, the convergence of adequately powered samples coupled with cutting-edge statistical and bioinformatics tools sets the scene for novel genetic and biological mechanisms underlying GCA to be made that might be amenable to novel therapeutic strategies.

Here, we jointly analysed the two largest GWAS of cognition to date^15,16^. In doing so, we harmonized the independent and/or convergent genome-wide signals associated with GCA across these studies at the levels of: individual variants, broader genomic regions of loci in linkage disequilibrium (LD), specific protein-coding loci/genes, and functional biological pathways. We also employed novel analytical methods not previously employed in cognitive GWAS studies to determine the direction of causality between GWAS hits for GCA and genetically correlated phenotypes. Large brain-based transcriptomic databases were then utilized to determine the biological underpinnings of the most credible and actionable cognitive GWAS to identify novel nootropic drug targets.

## Methods

Study workflow is detailed in Figure 1. As can be seen, data analyses comprised several stages. The core analysis combined summary statistics of the two largest GWAS of cognition to date^15,16^. Savage et al (N = 269,867) analysed 9,395,118 single nucleotide polymorphisms (SNPs) for association to intelligence, and Davies et al.^16^ (N = 283,531) analysed 12,871,771 SNPs in relation to the somewhat broader general cognition phenotype. The latter set of summary statistics reported by Davies et al.^16^ was reduced from the original N = 300,486 due to data access limitations. It is important to note that the two studies had a relatively large degree of sample overlap; while statistical inflation of the combined results is well-controlled by MTAG, the relative increase in power available is more modest than would be expected for a meta-analysis of fully independent samples.

**Figure 1.**
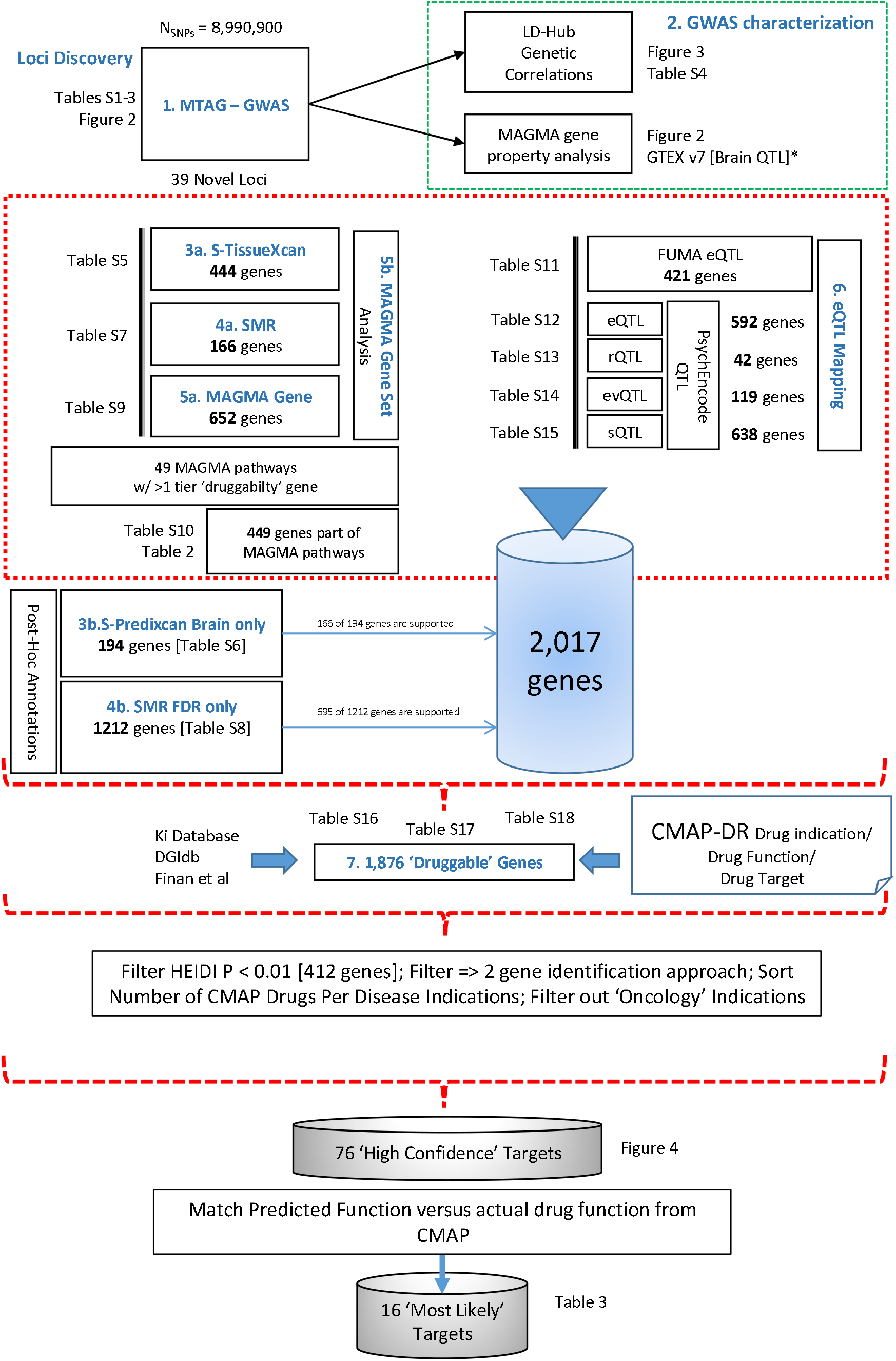
Workflow for GWAS, Gene Identification Approaches, and Drug Database Annotations

### 1. Loci Discovery: MTAG-GWAS

We meta-analysed both sets of summary statistics using the Multi-Trait Analysis of GWAS (MTAG^31^ v1.08). MTAG adjusts for sample overlaps based on LD score regression. It is notable based on reported sample sizes, that approximately 89% of samples between Davies et al.^16^ and Savage et al.^15^ As part of the MTAG workflow, alleles in both sets of summary statistics were aligned against the 1000 genomes phase 3 version 5a reference panel^54^. We set filters for sample size N > 10,000, and variations with minor allele frequency > 0.001. To obtain a single output from MTAG, we set covariance of both phenotypes to 1 and equivalent heritability across both phenotypes. This would approximate a fixed effect inverse variance meta-analysis but adjusting for sample overlaps across summary statistics input. We also carried out FDR Inverse Quantile Transformation analysis to account for potential Winner’s Curse^32^. Following MTAG, significant loci were identified as follows: First, independent lead SNPs were identified using the clumping function in FUMA v1.3.5^33^ with default parameters of *R*^2^ < 0.1, 250 kb window based on the 1000 Genomes Project Phase 3 European ancestry LD reference panel^34^. Independent loci were then identified by taking SNPs that are within an LD of *R*^2^ > 0.6 of the lead SNP. Finally, loci within 250kb of each other were merged into a single locus. GWAS significant threshold was set to *P* < 5*e* — 8, this was also the threshold used by FUMA to identify the lead SNPs with P-values less than or equal to the GWAS significant threshold. Candidate SNPs in LD with the significant independent SNP were selected based on the secondary *P* < 0.05 threshold. The minor allele frequency threshold for SNPs to be included in annotation and prioritization MAF > 0.01. We applied the default value of 10 kb for positional mapping of SNPs to genes, or functional consequences.

### 2. Genome-wide characterization

We then carried out phenome-wide genetic correlation analysis using LD-hub^35^ (v1.9.3) to determine and visualize the relationship between cognition and other psychiatric and physical traits. In addition, to confirm that our genetic discoveries reflected brain-based biological traits underlying cognitive performance, a gene property analysis was used to screen gene-expression and localization in CNS tissue vs. all other biological tissue as implemented in MAGMA^36^ utilizing GTEx v7 (http://www.gtexportal.org/home/datasets) tissues.

### 3. S-PrediXcan/S-TissueXcan Transcriptome-Wide analysis of gene expression

To expand our analysis from SNPs/loci and to identify putative causal genes, several methods were employed (See Workflow - Figure 1). First, genetically regulated gene expression was imputed for MTAG meta-analysis using tissue models from GTExv7, which contains 48 different tissue types across 30 general tissue categories. The summary statistics from this meta-analysis were entered into the S-PrediXcan (Web app 18 Apr 2019) framework (https://cloud.hakyimlab.org/). S-PrediXcan computes gene-based associations where genetic effects on phenotypes are mediated through gene expression^37^ (see also http://predictdb.org/). Next, we utilized S-TissueXcan to exploit the gene expression-mediated associations shared across multiple tissues to enhance power for gene identification. We combined all S-PrediXcan results based on individual tissue types in GTEx v7 using S-TissueXcan. Gene-based p-value is computed via an omnibus test, which are then Bonferroni corrected. Both S-TissueXcan processing and post-processing pipelines are available online (http://cloud.hakyimlab.org/). We paid special attention to S-PrediXcan’s brain tissue annotations in later stages of the analysis, extracting genes that are significantly associated with cognition after Bonferroni correction for each brain tissue within the GTEX database to lend post-hoc support for the gene-identification approaches.

### 4. Summary Statistics Based Mendelian Randomization (SMR and HEIDI)

As a more conservative approach to transcriptomic-based gene identification, we utilized SMR (Summary-based Mendelian Randomization) and HEIDI (Heterogeneity in Dependent Instruments) tests^30^ (v1.02). SMR uses a Mendelian Randomization (MR) approach where one or multiple SNPs could be used as instruments to identify gene expression effects on a given trait with estimated SNP-gene expression and SNP-phenotype effects. At the same time, the HEIDI test identifies SNP-gene expression effects and SNP-phenotype effects that are correlated with each other through LD rather than biologically related via pleiotropy or a causal pathway. We prioritized GWAS-identified genes for followup functional studies by including only genes with biologically related expression and phenotype effects (i.e., excluding genes with P_HEIDI_ < 0.01). For the SMR analyses, we utilized multiple transcriptomic reference datasets: (i) GTEx-brain eQTL data with estimated effective sample size (N) of 233, which includes an eQTL based meta-analysis of 10 brain regions from the GTEx, while correcting for sample overlap^24,25^; (ii) Brain-eMeta eQTL data with estimated effective N = 1,194, which includes an eQTL based meta-analysis of GTEx-brain, CommonMind Consortium, and xQTLServer (ROSMAP) datasets^24^; (iii) the PsychENCODE prefrontal cortex eQTL data (N = 1,866). Two sets of brain-based eQTL were generated from the PsychENCODE data based on earlier reports: (a) eQTL corrected for 50 Probabilistic Estimation of Expression Residuals (PEER), where only SNPs with expression association FDR <0.05 were included^28^ and (b) eQTL corrected for 100 Hidden Covariates with Prior knowledge (HCP) were included as covariates^29^. For brain-based eQTL datasets utilized by SMR and HEIDI, only SNPs within 1Mb of each probe were included as a proxy for *cis*-acting eQTL (See https://cnsgenomics.com/software/smr/#Overview). Both SMR and HEIDI p-values were Bonferroni corrected for multiple testing in 15,302 genes. Due to SMR’s more conservative estimation of p-values, we also performed Benjamini-Hochberg false discovery rate (FDR) correction for SMR genes, subsequent to the primary analysis.

### 5. MAGMA Gene- and Gene Set-based association analysis

MAGMA^36^ (v1.07) gene-based association tests were carried out as part of the FUMA pipeline. The MAGMA gene-based test combines individual SNP p-values in a pre-defined gene region into a gene-based p-value by calculating the mean chi-square statistics accounting for LD between SNPs and correcting for gene size. LD between SNPs within the genes is estimated based on the 1000 genomes phase 3 European ancestry panel. MAGMA competitive pathway analysis was conducted with results emerging from earlier MAGMA gene-based, S-PrediXcan/S-TissueXcan, and SMR/HEIDI analyses. Gene sets that were tested included custom-curated neurodevelopmental and other brain-related gene sets that had gone through stringent quality control in a study originally designed to interrogate rare variants in schizophrenia^38^. In the latter, pathways with more than 100 genes from Gene Ontology (release 146; June 22, 2015 release), KEGG (July 1, 2011 release), PANTHER (May 18, 2015 release), REACTOME (March 23, 2015 release), DECIPHER Developmental Disorder Genotype-Phenotype (DDG2P) database (April 13, 2015 release) and the Molecular Signatures Database (MSigDB) hallmark processes (version 4, March 26, 2015 release) were initially included. Additional gene sets were selected based on risk for schizophrenia and neurodevelopmental disorders, including those reported for schizophrenia rare variants^39^ (translational targets of FMRP^40,41^, components of the post-synaptic density^39,42^ ion channel proteins^39^, components of the ARC, mGluR5, and NMDAR complexes^39^, proteins at cortical inhibitory synapses^43,44^, targets of mir-137^39^, and genes near schizophrenia common risk loci^39,45^) and autism risk (These include: (1)targets of *CHD8*^46–48^ (2) splice targets of RBFOX^48–50^, (3) hippocampal gene expression networks^51^, (4) neuronal gene lists from the Gene2cognition database [http://www.genes2cognition.org]^48^, as well as (5) loss of function intolerant genes (pLI > 0.9 from the ExAC v0.3.1 pLI metric), (6) ASD exomes risk genes for FDR < 10% and 30%, and (7) ASD/developmental disorder *de novo* genes hit by a LoF or a LoF/missense *de novo* variant^52,53^). Brain-tissue expression genesets included the Brainspan RNA-seq dataset^54^ and the GTEx v7 dataset^25^. We report significant gene sets that were associated with GCA to identify biological pathways putatively associated with GCA. Moreover, we use this information to further interrogate genes within these pathways with protein products that may serve as druggable targets but which failed to attain genomewide significance on their own. As such, we extracted nominally significant (p<.05) genes within the significant gene sets for further drug target annotations; this threshold was selected to strike a balance between potential false positive and false negative associations within gene sets that had already demonstrated association signal to GCA.

### 6. Brain-based eQTL mapping

eQTL mapping was carried out as part of the FUMA pipeline. Brain eQTL annotations were utilized for eQTL mapping. Databases used for eQTL mapping include: (i) BRAINEAC (http://www.braineac.org). A total of 134 neuropathologically confirmed control individuals of European descent from UK Brain Expression Consortium were included in the BRAINEAC data. All eQTLs with nominal p-value < 0.05 were identified in the cerebellar cortex, frontal cortex, hippocampus, inferior olivary nucleus, occipital cortex, putamen, substantia nigra, temporal cortex, thalamus, and white matter regions and based on averaged expression across all of them. (ii) GTEX v7: For the purpose of eQTL mapping analysis, we chose brain tissue expression from GTEX v7 and defined significant eQTLs as FDR (gene q-value) < 0.05. The gene FDR is pre-calculated by GTEx and every gene-tissue pair has a defined p-value threshold for eQTLs based on permutation. (iii) xQTLServer (http://mostafavilab.stat.ubc.ca/xqtl/): Expression of dorsolateral prefrontal cortex from 494 samples, (iv) Brain expression from 467 Caucasian samples available at the CommonMind Consortium (https://www.synapse.org//#!Synapse:syn5585484). Publicly available eOTLs from CMC are binned by FDR into four groups: <0.2, <0.1, <0.05 and <0.01.

Finally, we mapped several novel forms of molecular quantitative trait loci (QTL). These novel Q,TL·s include expression variation, splicing, and translation using post-mortem prefrontal cortex tissue data from the PsychENCODE/BrainGVEX project. In these samples, gene transcription and translation activities were assayed by RNA-sequencing (N=416) and ribosome profiling (N=192); Annotation data was available for novel QTLs with SNPs at *P_MTAG_* < 1 *x* 10^-5^: (i) gene expression variation QTL (evQTL) analysis tests for genetic loci that influence variance of expression level, using Bartlett’s test^55^ on the RNA-seq data; (ii) splicing QTL (sQTL) analysis captures the effects of genetic variations on RNA splicing, using leafcutter^56^ on RNA-seq data (iii) ribosome occupancy QTL (rQTL) analysis identifies genetic variations that influence translation-related ribosome occupancy using Ribo-seq data; prior reports suggest that differences between transcription and translation QTLs may yield novel biological insights beyond standard eQTLs alone^57,58^. We focused on the cis-QTL by testing SNPs within 1 Mb of genes for the three molecular phenotypes. Significant QTL association was defined by FDR p value < 0.05. For this analysis, we carried out mean-variance QTL mapping, using on the double generalized linear model approach discussed in detail elsewhere^55,59^. In prior simulations, mean-variance approach to QTL mapping and associated permutation procedures have shown to be robust in reliably identifying QTL in face of variance heterogeneity. For comparison purposes, we also performed standard eQTL mapping on this dataset.

### 7. “Druggable” Gene Annotations

We identified a set of “druggable” gene targets derived from the Drug-Gene Interaction database (DGIdb v.2), Psychoactive Drug Screening Database K_i_DB, and a recent review on “druggability”^60^. The “Druggable genome” as previously identified by Finan et al.,^60^ includes 4,465 gene targets and is annotated into 3 Tiers based on “druggability” levels: (i) Tier 1 gene targets are those derived from FDA-approved compounds, or from compounds that are presently studied in clinical trials; (ii) Tier 2 gene targets include genes with high sequence similarity to Tier 1 proteins, or those that are targeted by small drug-like molecules; (iii) Tier 3 gene targets code for secreted and extracellular proteins, which also belong to “druggable” gene families. DGIdb v.2 integrates drug-gene interactions from 15 databases, including DrugBank and ChEMBL. The data is directly available as drug-gene pairs; the K_i_DB provides K_i_ values for drug/target pairs and is particularly relevant for psychoactive drugs. Using filtering criterion previously reported by Gaspar and Breen^21^ (K_i_DB: “With non-empty K_i_ field”, “Only Human”, “K_i_ not superior or inferior to a value”, “With molecule name”, “With gene-name”, “Unique pairs”, “With range pK_i_ > 2”; DGIdb: “Number of unique gene-sets > 2”), we identified and updated 2,567 potential gene targets from the chemoinformatic databases. For further gene-target annotations, we took the intersection between genes extracted from the chemoinformatic database, and those reported in Finan et al.,^60^. This resulted in 1,876 genes for further annotations. At the final stage of the analysis we annotated high confidence genes using the Broad Institute Connectivity Map, Drug Re-purposing Database^61^ which provides more in-depth details such as drug names, mechanism of action, and drug indications.

## Results

### 1. Loci Discovery: MTAG-GWAS

We carried out MTAG meta-analysis^31^ on the two largest GWAS of cognition. Median value of Z-scores in Savage et al.^9^ was −0.001, mean *χ*^2^ = 1.688, and genomic inflation was *λ_GC_* = 1.456; 13,173 SNPs passed genome-wide significant thresholds (*p* < 5 *x* 10^-8^) and the LD-score regression (LDSC)^62,63^ intercept was 1.051. Median value of Z-scores in Davies et al.^16^ was 0.0, mean *χ*^2^ = 1.455, and genomic inflation was *λ_GC_* = 1.307; 11,244 SNPs passed genome-wide significant thresholds, and the LDSC-intercept was 1.038. A total of 8,990,900 SNPs was present in both sets of summary statistics were extracted for use in the MTAG meta-analysis. Since both sets of GWAS summary statistics indexed GCA, we constrained MTAG analysis to give a single output; heritability and genetic covariance for phenotypes in either set of summary statistics was set to be the same. A grid search for maximum potential for false discovery using MTAG revealed very low probability of false positives (*max_fdr_* = 4.51 *X* 10^-7^). The resulting mean chi-square values were as follows: mean 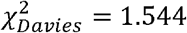, mean 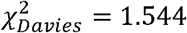, and mean 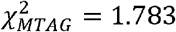. The average projected GWAS equivalent sample size increase after MTAG analysis for Savage et al.^9^ was *N_Savage_* = 338,737 and for Davies et al.^10^ *Noavies* = 408,498, (approximate estimated sample size ~ 373,617) which shows substantial power improvement.

Clumping procedures were carried out on 8,990,900 SNPs (See Methods). SNPs were extracted based on *R*^2^ > 0.6 within each independent LD-block; we identified SNPs within 250kb of each other as an independent locus. These loci definitions were merged with previously reported loci by Savage et al.^15^ and Davies et al.^16^. A total of 304 genomic loci were identified as potentially associated with GCA using this method. Of these, 241 loci were GWAS significant for the MTAG analysis (Figure 2), while 214 loci and 124 loci were GWAS significant for Savage et al.^15^, and Davies et al.^16^ respectively. It should be noted that 17 loci were not significant in any sets of summary statistics, likely due to the sample size reduction in the Davies et al.^16^. We also note that 38 loci reported as significant in Savage et al.^15^ and 8 loci in Davies et al.^16^ were no longer significant in the MTAG analysis (Supplementary Table 2; Figure 2). Winner’s curse analysis suggested that these loci were likely false positives in the original studies (Supplementary Table 3). A total of 39 of these were novel and not found in the input GWAS. We then looked up reports that have used multi-trait strategies to enhance power for GCA^17,20^ and found that of the 39 loci, 8 loci were also reported by Hill et al.^17^, 1 locus was reported by Lam et al.^20^ and 1 locus was reported by both of these studies.

**Figure 2.**
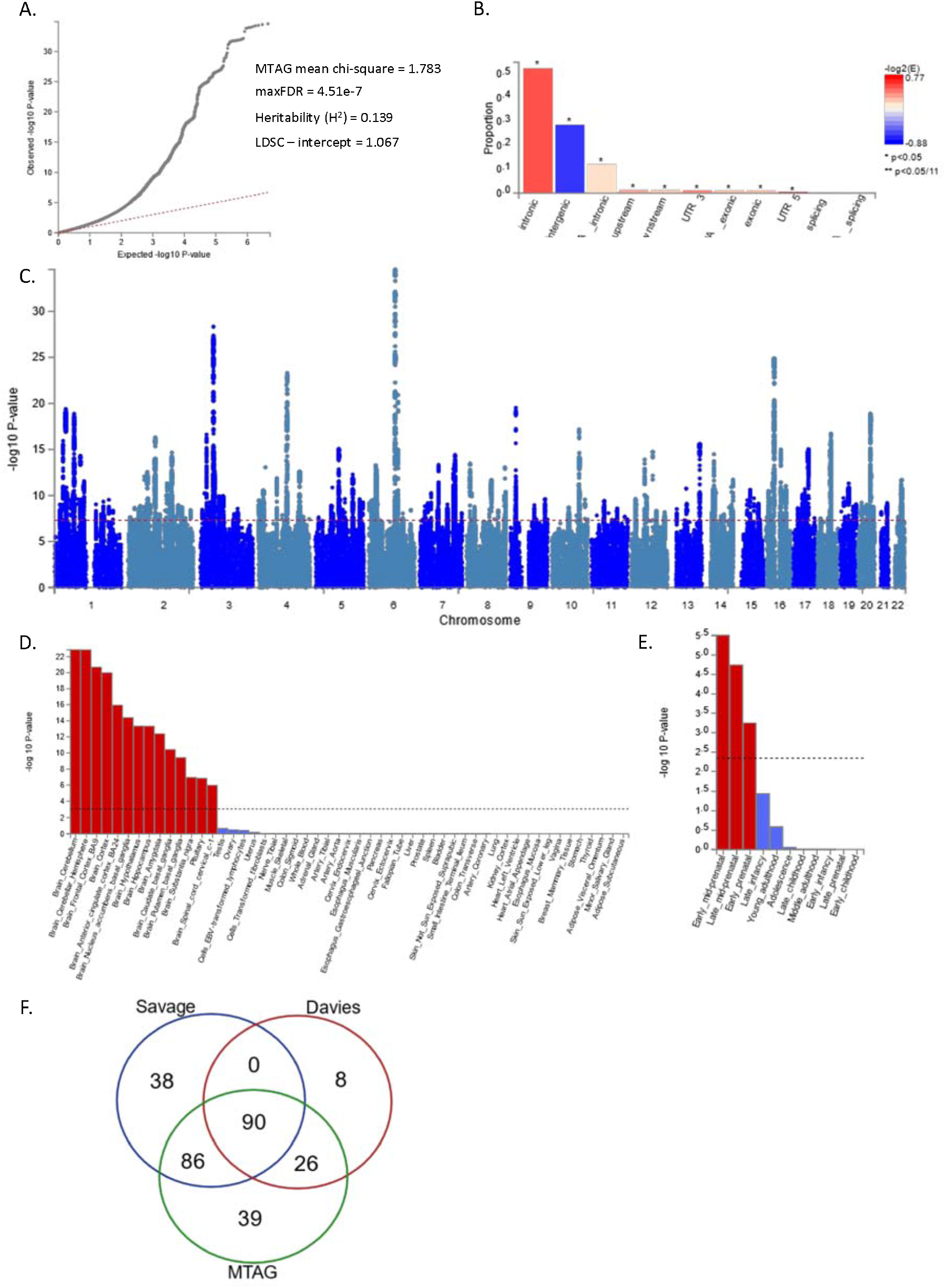
GWAS association plots for Cognitive MTAG. A. QQ-plot B. SNP annotation plot C. MAGMA gene property analysis for overall GTEXv7 D. MAGMA gene property analysis using BrainSpan E. Venn Diagram showing loci overlap

### 2. Genome-wide characterization

Genetic correlations were conducted between GCA and 855 phenotypes from LD-hub^28^ and UK Biobank. MTAG summary statistics were merged and aligned with HapMapЗ SNPs without the MHC region for genetic correlation analysis (1,190,946 SNPs remained). The reduced set of SNPs had a median Z-score of −0.001, mean *χ*^2^ = 2.120, LDSC-intercept = 1.067 and *h*^2^ = 0.139. Of these, 297 phenotypes showed significant genetic correlation with cognition at *P_Bonferroni_* < 0.05. Consistent with prior reports^10,11,17,64^, traits genetically correlated with GCA, included education, reproduction, longevity, personality, smoking behavior, anthropometric, brain volume, psychiatric, dementias, lung function, sleep, glycemic, autoimmune, cardio-metabolic, cancer and several ICD-10 medical phenotypes. Several novel traits that have not been previously reported to be genetically correlated with GCA are displayed in Figure 3. Full results are reported in Supplementary Table 4.

**Figure 3.**
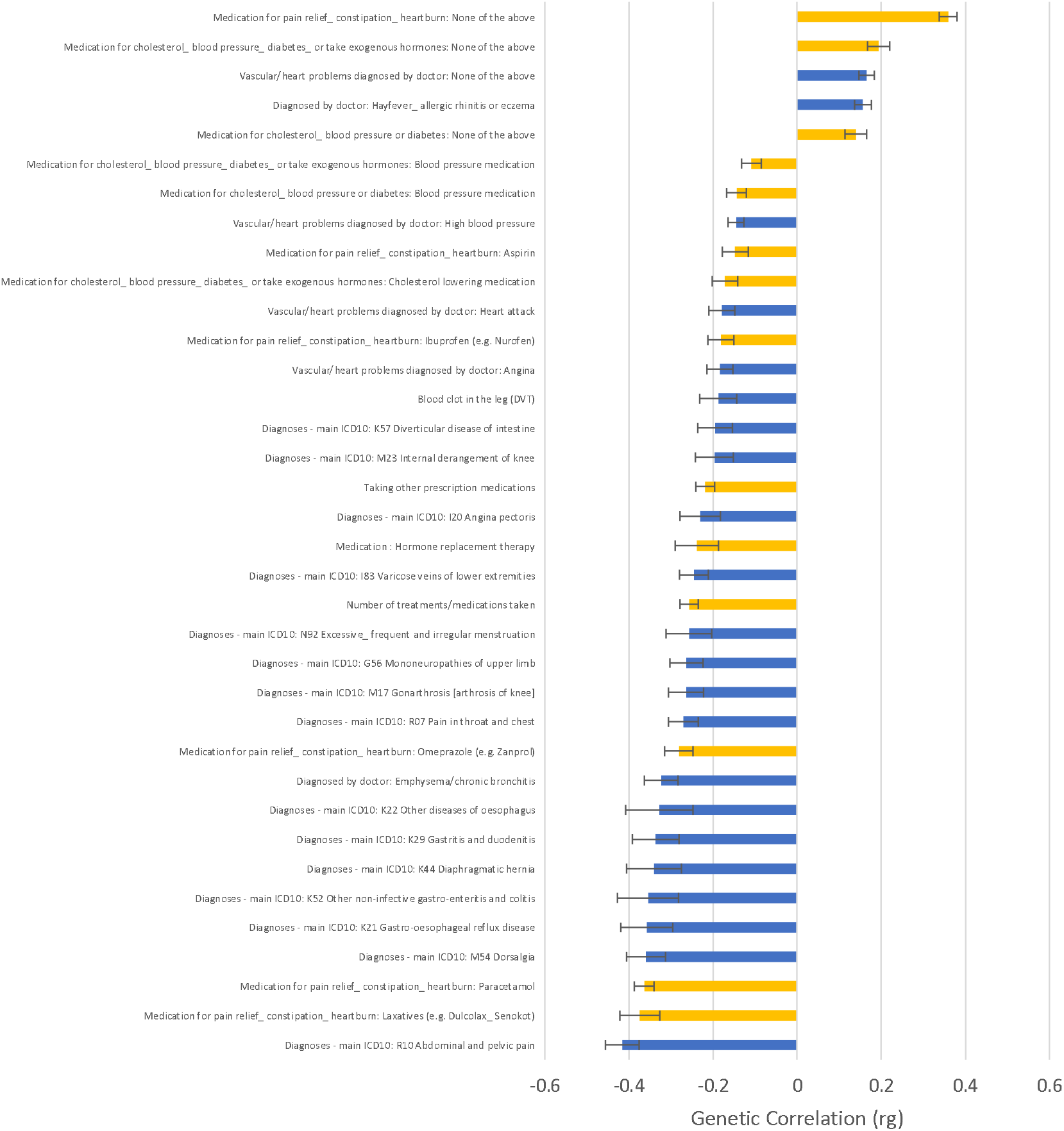
Genetic Correlations for UK Biobank ICD-10 and Medication phenotypes. Error bars denote standard errors. Yellow bars denote medication phenotypes. Blue bars denote ICD10 phenotypes.

#### SprediXcan/S-TissueXcan, SMR &HEIDI, MAGMA Gene Set Analysis and Brain-based eQTL mapping

### 3. SprediXcan/S-TissueXcan

As described (Figure 1), we conducted a variety of complementary transcriptomics analyses, in order to convert SNP/locus results into directional, biologically interpretable, gene effects on GCA. S-TissueXcan^37^ analysis carried out in all 48 GTEXv7 tissues yielded 444 significant genes after Bonferroni correction (Supplementary Table 5). Of these, 194 genes were significant in one or more S-PrediXcan brain tissue annotations (Supplementary Table 6).

### 4. SMR &HEIDI

Using SMR, we were able to identify 166 genes that were genome-wide significant, where gene expression levels were contributing to variance of GCA (Supplementary Table 7). As discussed previously, SMR analysis tended to be more conservative than other gene identification methodologies^37^, hence we computed using the Benjamini and Hochberg method, FDR corrected p-values for nominally significant genes (*P_SMR_* < 0.05) The second approach yielded 1212 genes for follow up in the later gene annotation (Supplementary Table 8). Importantly, there were 412 genes associated with linkage *P_HEIDI_* <0.01 and these were excluded from subsequent “druggability” analysis.

### 5. MAGMA Gene Set Analysis

MAGMA gene-based analysis revealed that 652 of 18,730 genes were significantly associated with GCA after Bonferroni correction (Supplementary Table 14). MAGMA pathway analyses were carried out using gene-based effect sizes from MAGMA gene-based analysis, SMR analysis (only using PsychENCODE results), and S-TissueXcan analysis. MAGMA significant pathways after Bonferroni correction are reported in Supplementary Table 15. Additionally, we annotated all genes within each significant MAGMA significant gene sets, with p-values from the other earlier gene identification approaches described above. Of 54 gene sets identified as significantly (following Bonferroni correction) associated with GCA, 49 included Tier 1 “druggable” gene targets (Supplementary Table 15; Table 2).

**Table 1.**
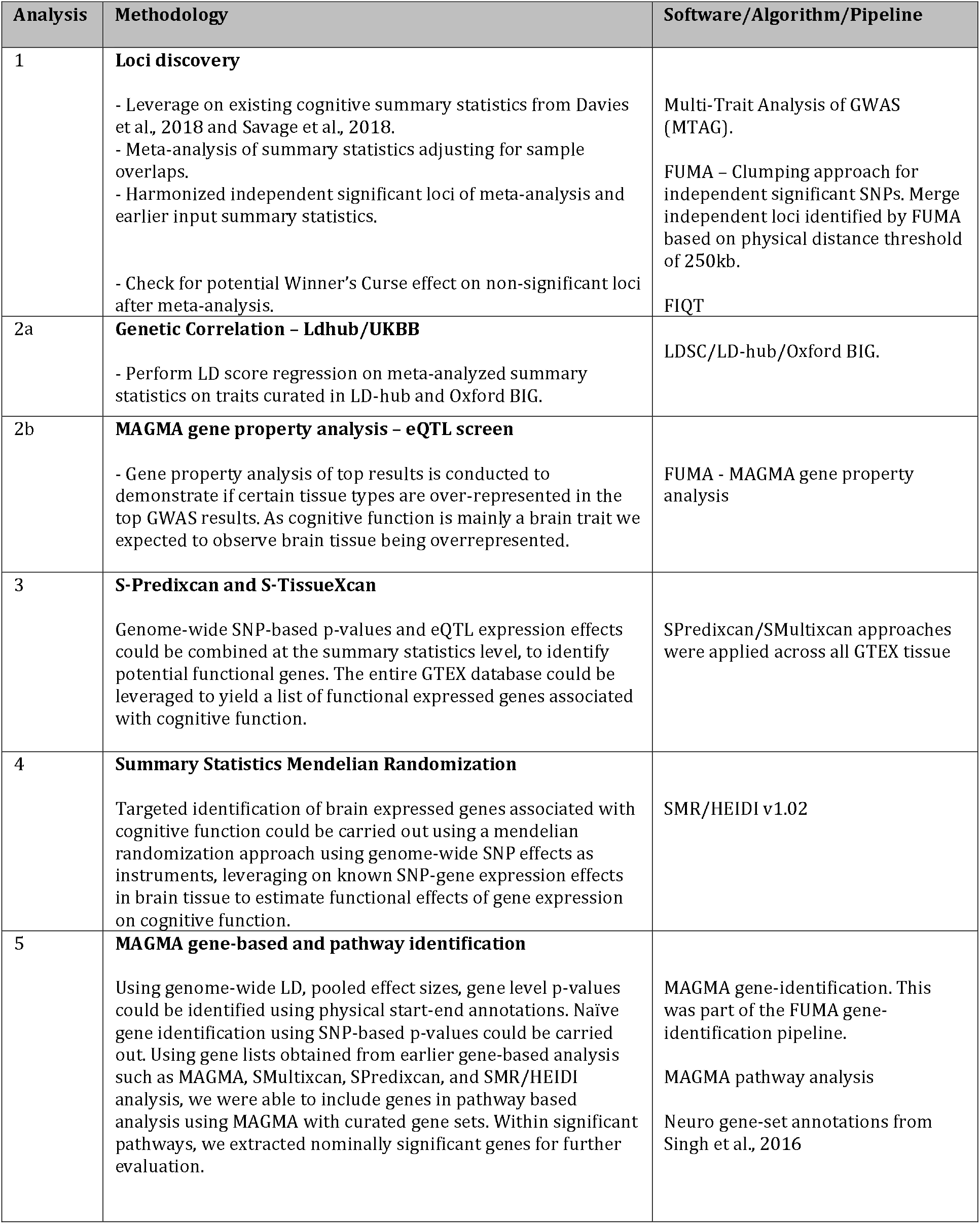
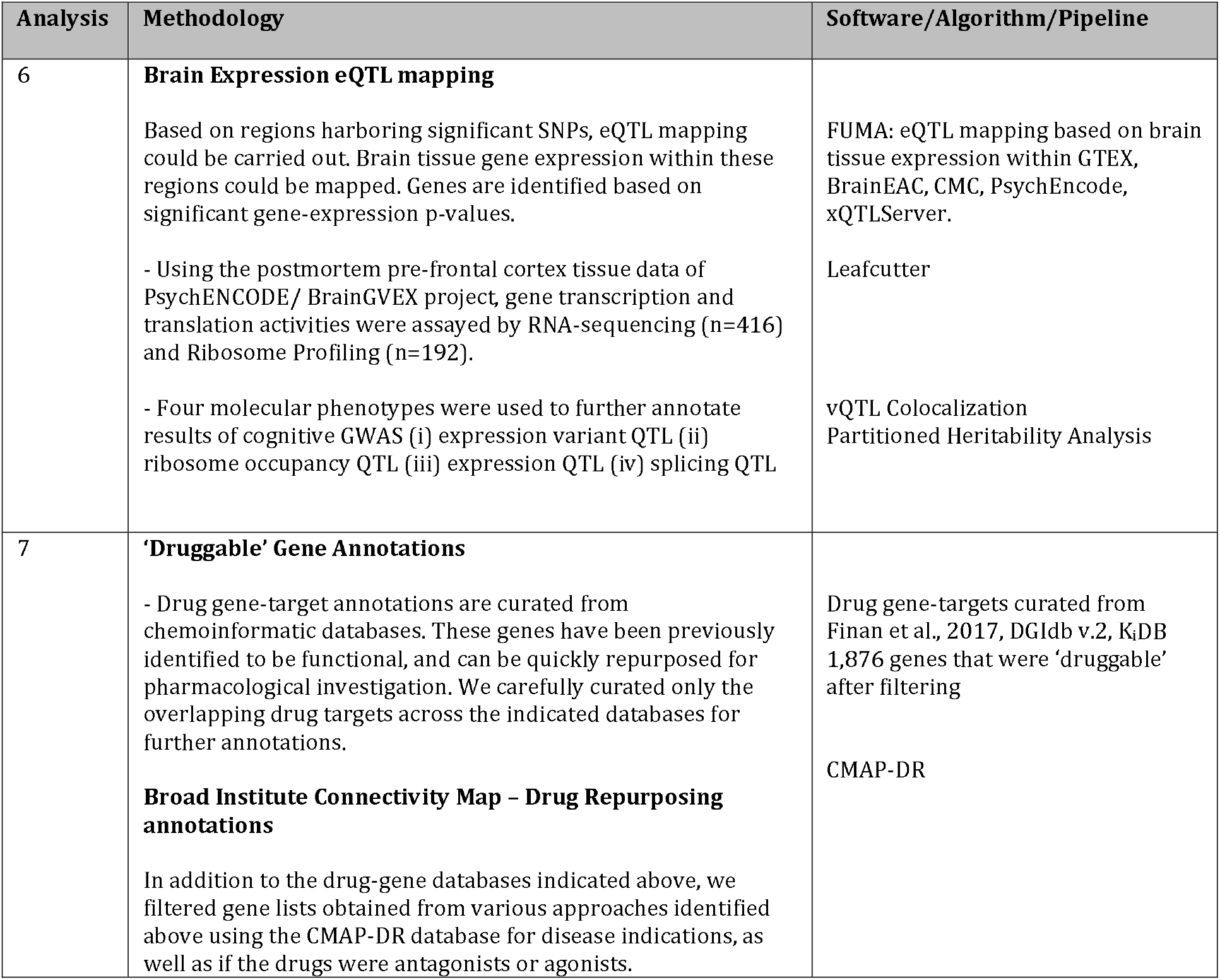
Methodological Overview and Analytical Approaches

| Analysis | Methodology | Software/Algorithm/Pipeline |
| --- | --- | --- |
| 1 | **Loci discovery**<br>- Leverage on existing cognitive summary statistics from Davies et al., 2018 and Savage et al., 2018.<br>- Meta-analysis of summary statistics adjusting for sample overlaps.<br>- Harmonized independent significant loci of meta-analysis and earlier input summary statistics.<br>- Check for potential Winner's Curse effect on non-significant loci after meta-analysis. | Multi-Trait Analysis of GWAS (MTAG).<br>FUMA – Clumping approach for independent significant SNPs. Merge independent loci identified by FUMA based on physical distance threshold of 250kb.<br>FIQT |
| 2a | **Genetic Correlation – Ldhub/UKBB**<br>- Perform LD score regression on meta-analyzed summary statistics on traits curated in LD-hub and Oxford BIG. | LDSC/LD-hub/Oxford BIG. |
| 2b | **MAGMA gene property analysis – eQTL screen**<br>- Gene property analysis of top results is conducted to demonstrate if certain tissue types are over-represented in the top GWAS results. As cognitive function is mainly a brain trait we expected to observe brain tissue being overrepresented. | FUMA - MAGMA gene property analysis |
| 3 | **S-Predixcan and S-TissueXcan**<br>Genome-wide SNP-based p-values and eQTL expression effects could be combined at the summary statistics level, to identify potential functional genes. The entire GTEx database could be leveraged to yield a list of functional expressed genes associated with cognitive function. | SPredixcan/SMultixcan approaches were applied across all GTEx tissue |
| 4 | **Summary Statistics Mendelian Randomization**<br>Targeted identification of brain expressed genes associated with cognitive function could be carried out using a mendelian randomization approach using genome-wide SNP effects as instruments, leveraging on known SNP-gene expression effects in brain tissue to estimate functional effects of gene expression on cognitive function. | SMR/HEIDI v1.02 |
| 5 | **MAGMA gene-based and pathway identification**<br>Using genome-wide LD, pooled effect sizes, gene level p-values could be identified using physical start-end annotations. Naïve gene identification using SNP-based p-values could be carried out. Using gene lists obtained from earlier gene-based analysis such as MAGMA, SMultixcan, SPredixcan, and SMR/HEIDI analysis, we were able to include genes in pathway based analysis using MAGMA with curated gene sets. Within significant pathways, we extracted nominally significant genes for further evaluation. | MAGMA gene-identification. This was part of the FUMA gene-identification pipeline.<br>MAGMA pathway analysis<br>Neuro gene-set annotations from Singh et al., 2016 |

Table 1 (cont'd). Methodological Overview and Analytic Approaches
| Analysis | Methodology | Software/Algorithm/Pipeline |
| --- | --- | --- |
| 6 | **Brain Expression eQTL mapping**<br>Based on regions harboring significant SNPs, eQTL mapping could be carried out. Brain tissue gene expression within these regions could be mapped. Genes are identified based on significant gene-expression p-values.<br>- Using the postmortem pre-frontal cortex tissue data of PsychENCODE/ BrainGVEX project, gene transcription and translation activities were assayed by RNA-sequencing (n=416) and Ribosome Profiling (n=192).<br>- Four molecular phenotypes were used to further annotate results of cognitive GWAS (i) expression variant QTL (ii) ribosome occupancy QTL (iii) expression QTL (iv) splicing QTL | FUMA: eQTL mapping based on brain tissue expression within GTEX, BrainEAC, CMC, PsychEncode, xQTLServer.<br>Leafcutter<br>vQTL Colocalization<br>Partitioned Heritability Analysis |
| 7 | **'Druggable' Gene Annotations**<br>- Drug gene-target annotations are curated from chemoinformatic databases. These genes have been previously identified to be functional, and can be quickly repurposed for pharmacological investigation. We carefully curated only the overlapping drug targets across the indicated databases for further annotations.<br>**Broad Institute Connectivity Map – Drug Repurposing annotations**<br>In addition to the drug-gene databases indicated above, we filtered gene lists obtained from various approaches identified above using the CMAP-DR database for disease indications, as well as if the drugs were antagonists or agonists. | Drug gene-targets curated from Finan et al., 2017, DGIdb v.2, K<sub>i</sub>DB 1,876 genes that were 'druggable' after filtering<br>CMAP-DR |

**Table 2a.**
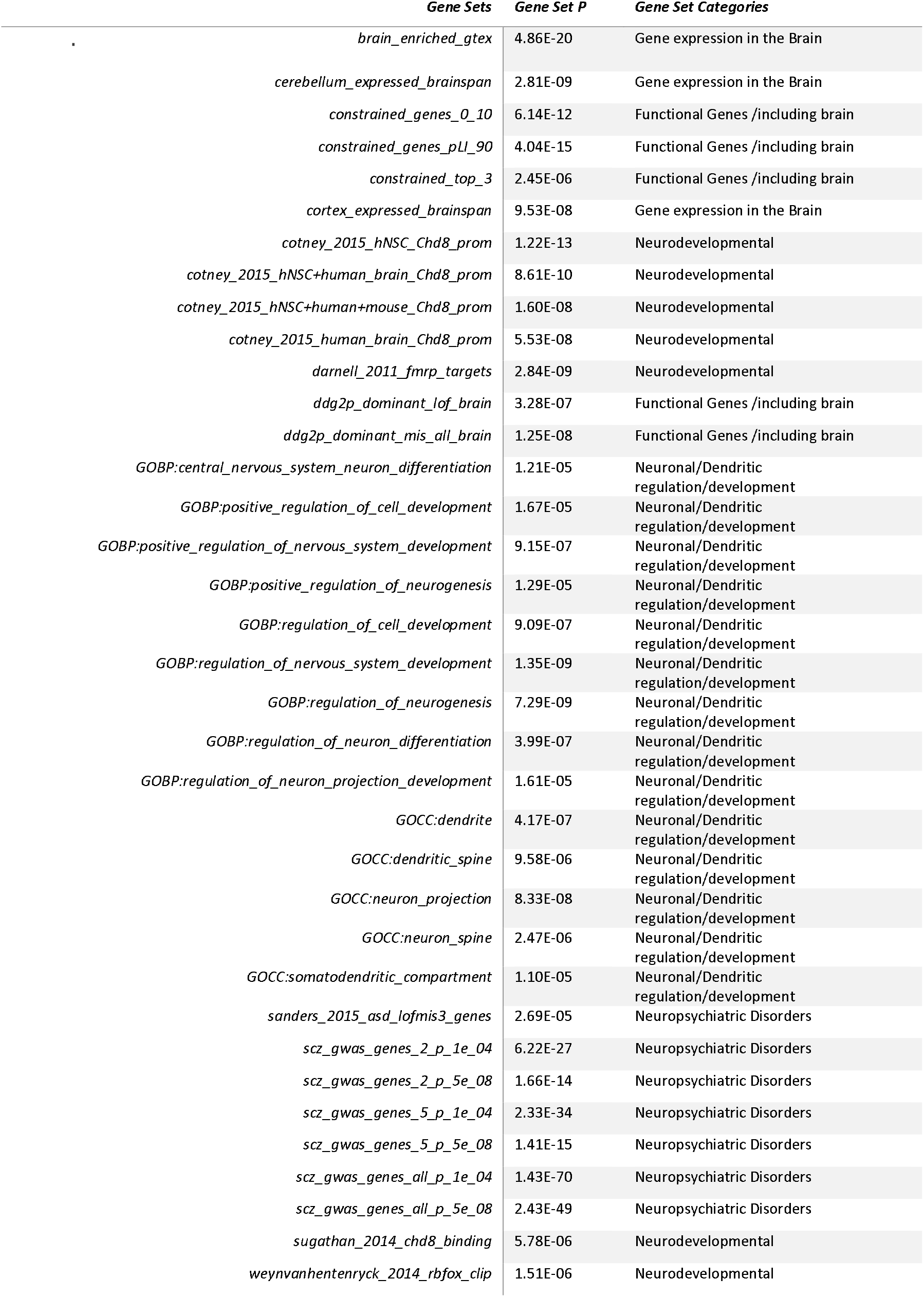
MAGMA Gene Sets Associated with Cognitive Function

**Table 2b.**
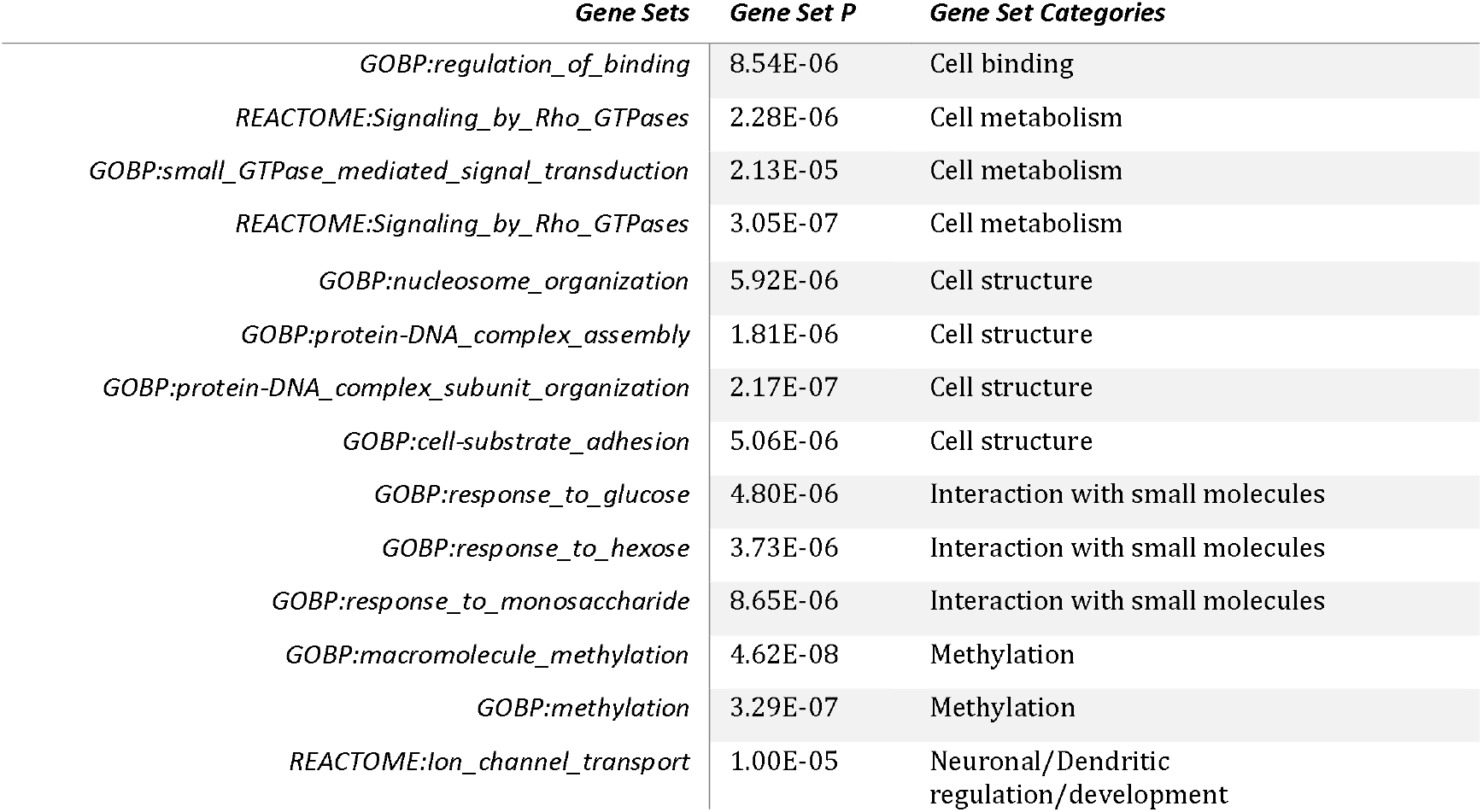
SMR Gene Sets Associated with Cognitive Function

| *Gene Sets* | *Gene Set P* | *Gene Set Categories* |
| --- | --- | --- |
| *GOBP:regulation_of_binding* | 8.54E-06 | Cell binding |
| *REACTOME:Signaling_by_Rho_GTPases* | 2.28E-06 | Cell metabolism |
| *GOBP:small_GTPase_mediated_signal_transduction* | 2.13E-05 | Cell metabolism |
| *REACTOME:Signaling_by_Rho_GTPases* | 3.05E-07 | Cell metabolism |
| *GOBP:nucleosome_organization* | 5.92E-06 | Cell structure |
| *GOBP:protein-DNA_complex_assembly* | 1.81E-06 | Cell structure |
| *GOBP:protein-DNA_complex_subunit_organization* | 2.17E-07 | Cell structure |
| *GOBP:cell-substrate_adhesion* | 5.06E-06 | Cell structure |
| *GOBP:response_to_glucose* | 4.80E-06 | Interaction with small molecules |
| *GOBP:response_to_hexose* | 3.73E-06 | Interaction with small molecules |
| *GOBP:response_to_monosaccharide* | 8.65E-06 | Interaction with small molecules |
| *GOBP:macromolecule_methylation* | 4.62E-08 | Methylation |
| *GOBP:methylation* | 3.29E-07 | Methylation |
| *REACTOME:ion_channel_transport* | 1.00E-05 | Neuronal/Dendritic regulation/development |

**Table 2c.** S-TissueXcan Gene Sets Associated with Cognitive Function

| *Gene Sets* | *Gene Set P* | *Gene Set Categories* |
| --- | --- | --- |
| *GOMF:actin_binding* | 6.44E-06 | Cell binding |
| *GOMF:chromatin_binding* | 4.52E-06 | Cell binding |
| *GOBP:cellular_macromolecular_complex_assembly* | 1.71E-07 | Cell structure |
| *GOBP:cellular_protein_complex_assembly* | 1.68E-07 | Cell structure |
| *GOBP:cellular_response_to_interferon-gamma* | 2.67E-06 | Interaction with small molecules |

A total of 449 genes were identified as part of significant MAGMA pathways. As shown in Table 2, several gene sets were identified to be significantly associated with GCA, based on the lists of significant genes derived from MAGMA, SMR, or S-TissueXcan. Not surprisingly, gene sets that have been associated with neuropsychiatric disorders such as schizophrenia and ASD were highly significant, consistent with significant genetic correlations between GCA and these disorders. Relatedly, gene sets related to neurodevelopmental processes implicated in schizophrenia and ASD, including the CHD8, FMRP, and RBFOX pathways, were also implicated in GCA^65^. Consistent with prior reports, a series of neuronal and dendritic development, differentiation, and regulation gene sets were associated with GCA^17^.

There were also several classes of gene sets emerging from our data that are novel with respect to GCA; notably, these results emerged in the context of the SMR and S-TissueXcan results, demonstrating the value of leveraging multiple approaches to post-GWAS gene identification. First, genes responsible for cellular response to small molecules such as sugars and cytokines appear to be implicated. Cell signal transductions mediated by small monomeric GTPases also appear to be relevant for GCA. In addition, genes sets underpinning cell structure and binding mechanisms, including adhesion, protein complexes, acting and chromatin binding were identified. It should be noted that gene sets representing methylation processes, DNA complex, and nucleosomes, while significantly associated with GCA, do not contain any genes that are targeted by known drugs, based upon our “druggability” criteria described in the Methods section above.

### 6. Brain-based eQTL mapping

eQTL mapping for gene expression across brain tissue indexed by the FUMA^33^ pipeline revealed 421 significantly expressed genes within GWAS significant regions (FDR corrected p-values; Supplementary Table 9). Additional mean variance QTL mapping of prefrontal cortex eQTL with GCA SNPs identified 638 genes with splicing activity, 42 genes implicated in ribosomal occupancy, 119 genes with expression variation levels, and 592 genes with eQTL (Supplementary Tables 10-13).

### 7. Identifying Drug-Gene Targets for Nootropic Re-purposing

A total of 2,017 genes (See Figure 1) were identified via one or more methods discussed in the previous sections. At this stage, we also merged additional post-hoc SMR analysis using FDR correction of p-values (N genes = 695), and the S-PrediXcan brain tissue eQTL analysis (N genes = 166) for further annotation. Filtering on the 1,876 “druggable” genes identified in the earlier step, genes with *P_HEIDI_* > 0.01, and genes that were identified by two or more gene-identification approaches, 91 genes were identified. We further annotated these genes with information from the Broad Institute CMAP Drug Repurposing Database^61^, drug indications for “Oncology” were filtered out mainly for drug delivery concerns; 76 “high-confidence” genes remained (Figure 4). These were annotated with eQTL directions (i.e., up-or down-regulation associated with higher GCA) for each gene. eQTL directions were obtained from earlier analysis, including brain-eQTLs from S-PrediXcan, SMR, PsychENCODE eQTL, RNA-seq Ribosomal and Splicing eQTL mapping, and overall S-TissueXcan GTEX eQTL analysis. Effect sizes that indicated up-regulation of the gene was denoted as “↑” and those that were down-regulated were denoted as “↓” (Supplementary Table 17). We predicted the “mechanism of action” from the overall eQTL direction if each gene might require either an “Agonist” or “Antagonist” to enhance GCA. This is achieved by taking the sum of eQTL directions across tissues (See Supplementary Table 17). If overall eQTL indicates up-regulation, it would be more likely require an agonist and vice-versa. We eliminated “Ambiguous” gene-targets which have an equal number of tissues that show up- and down-regulated gene expression.

**Figure 4.**
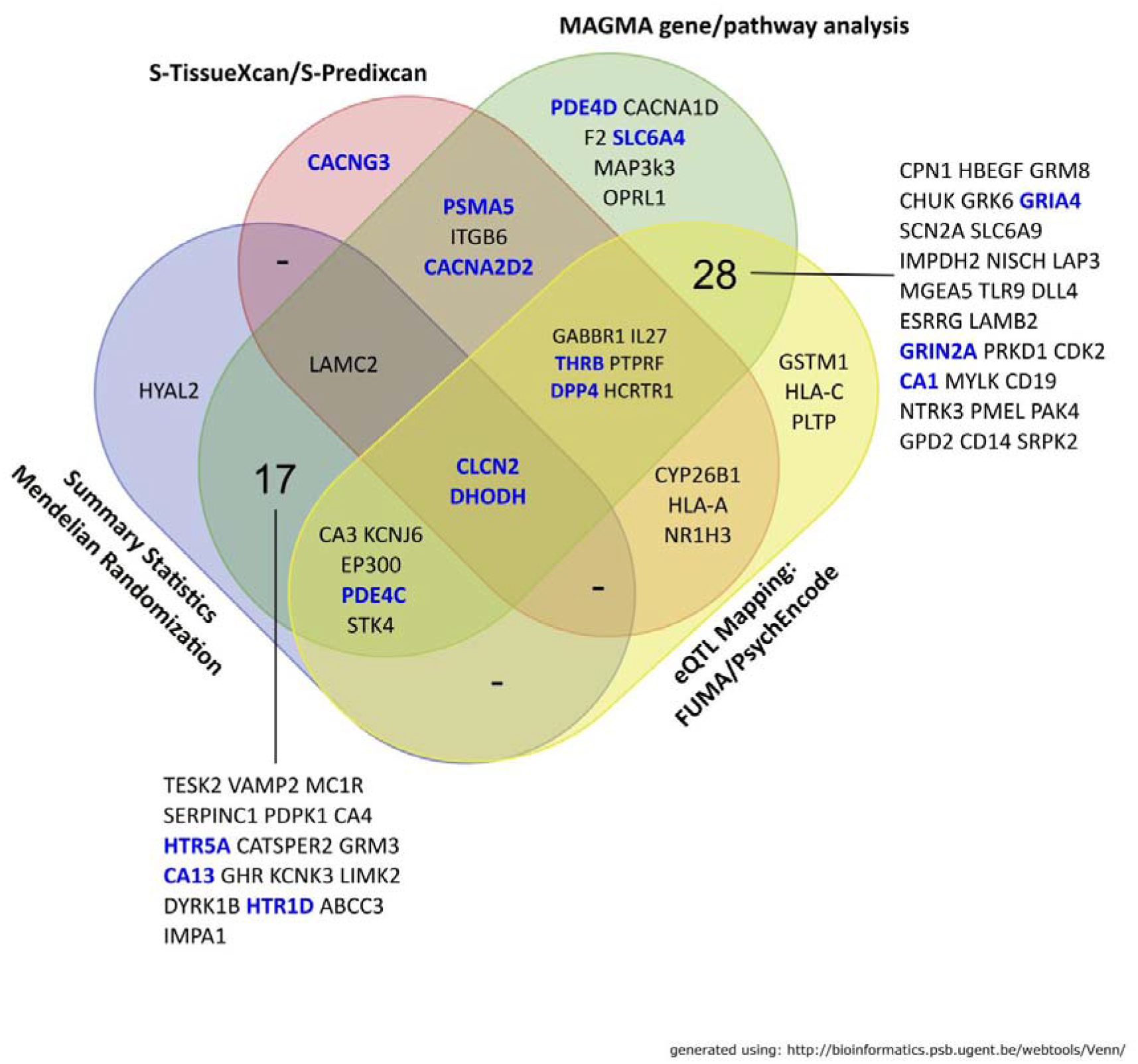
Venn Diagram of “High Confidence” Genes and Gene Identification Approaches. Genes highlighted in blue were deemed as most likely having gene targets that were suitable for nootropic re-purposing

CMAP Drug Re-purposing annotations which include drug names, mechanism of action (MOA), as well as drug indications were merged with the “high” confidence genes (Supplementary Table 18). We further filtered the high confidence genes based on the predicted and actual MOA. We were able to identify potential drugs related to the matched MOA, and their indications with various types of physical or psychiatric conditions are listed in Table 3. Notably, the relationship between some of the drug MOA and gene targets do not always appear to be direct. For example, adrenergic receptor agonists can indirectly activate calcium channels of which *CACNA2D2* is a constituent.

**Table 3.**
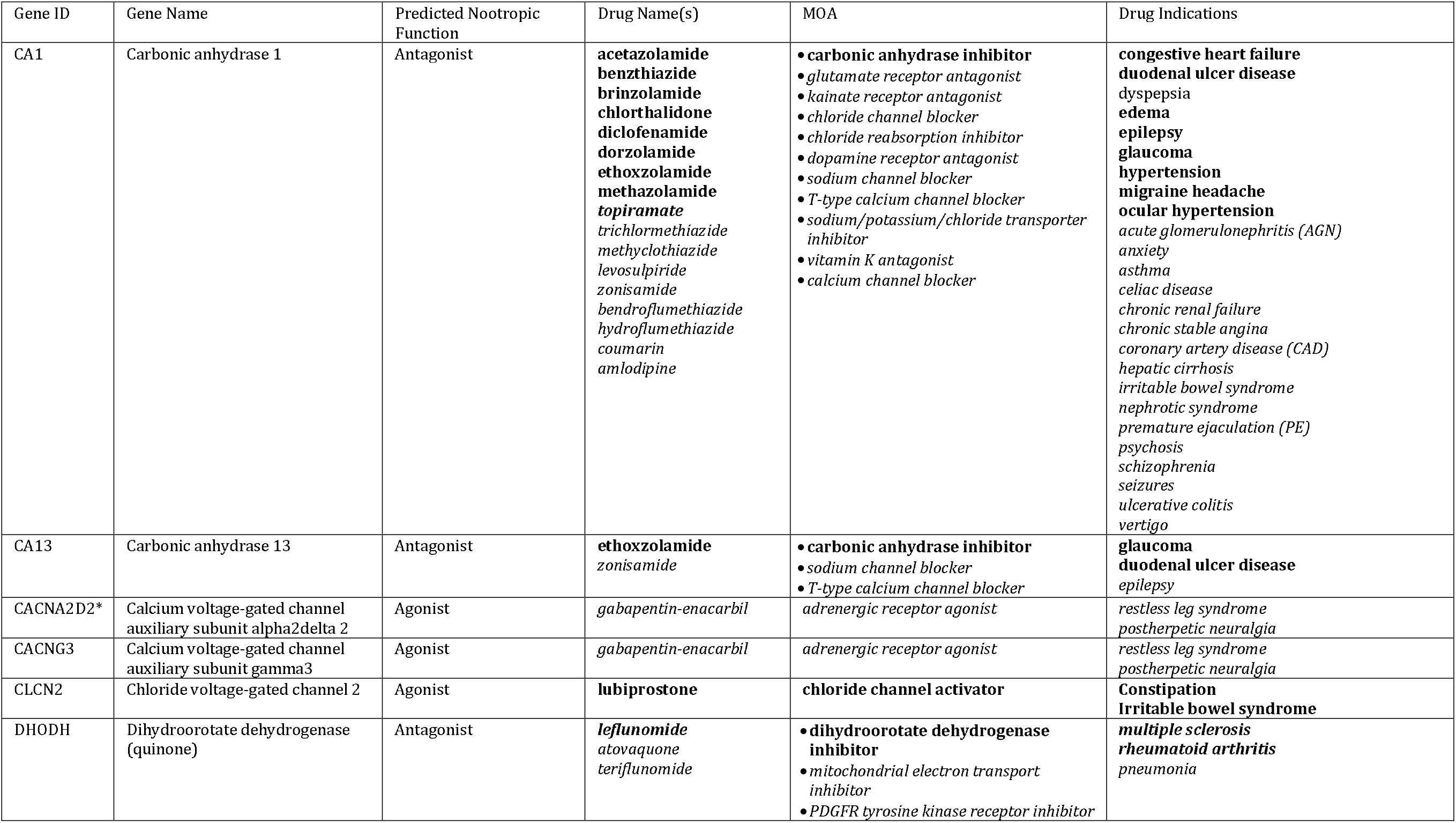
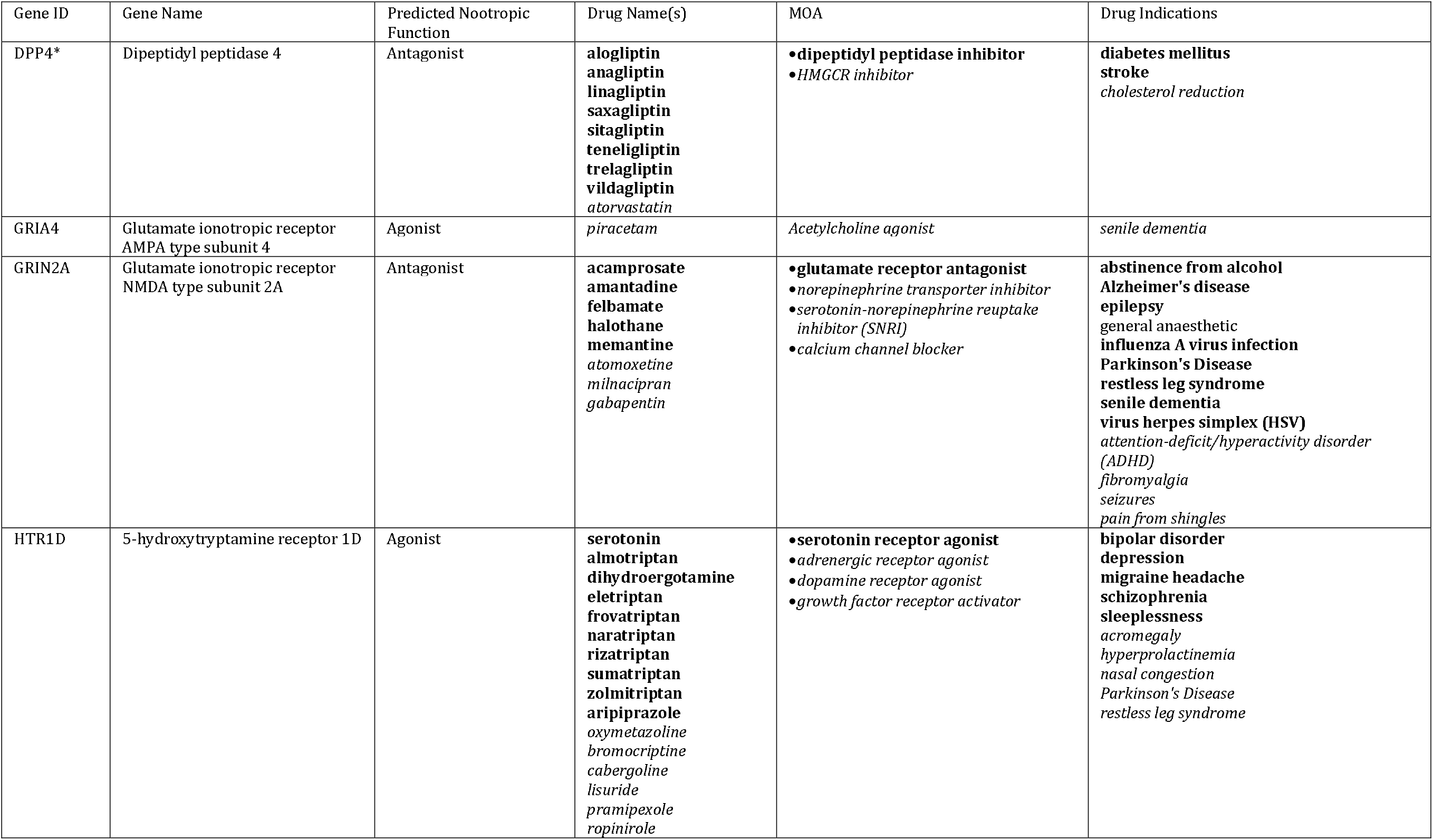
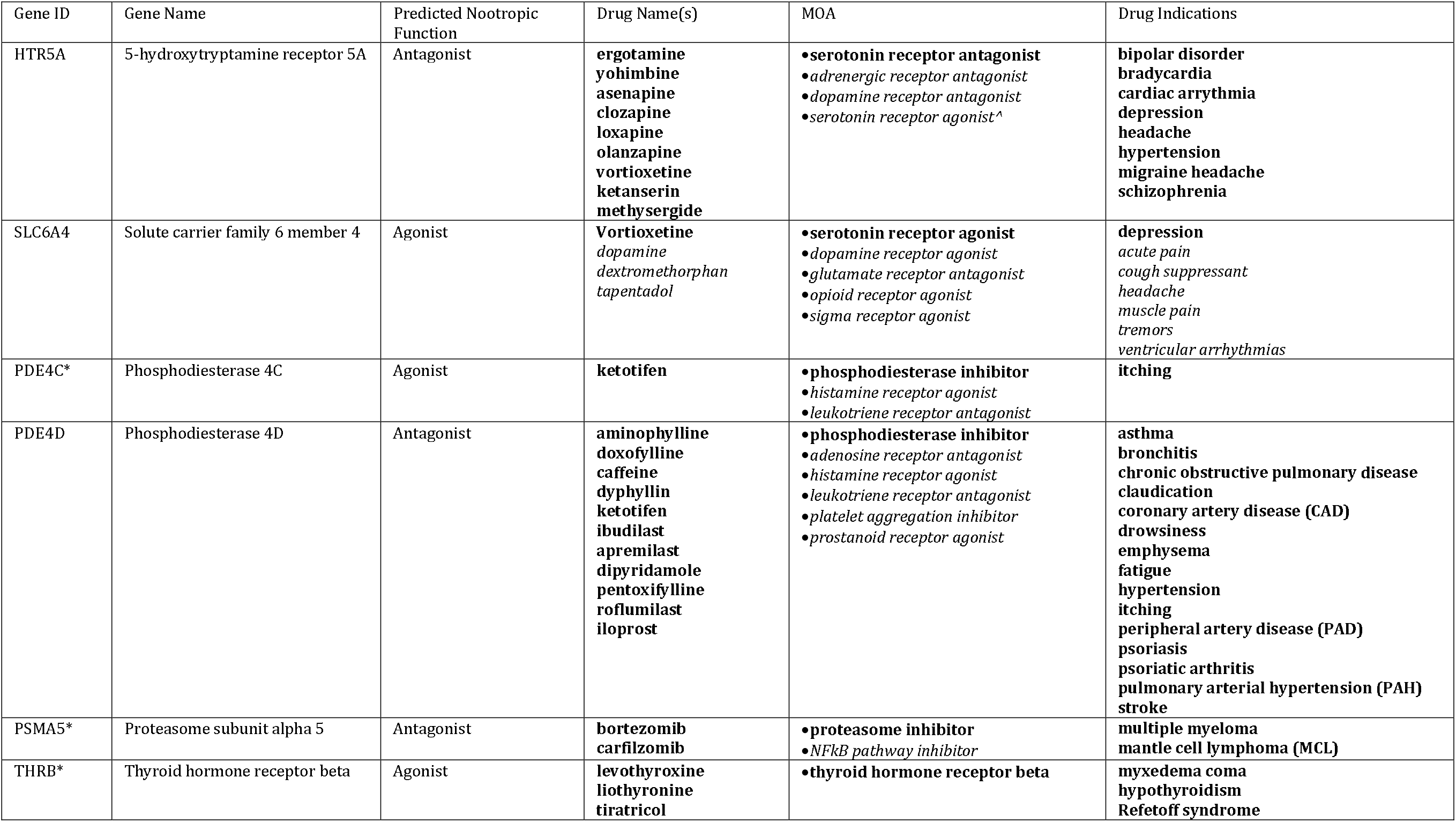
Prioritized Gene for Nootropic Re-purposing. *Note:* MOA: Mechanism of Action. Predicted Nootropic Function was obtained from gene expression association with general cognitive ability. MOA, Drug names and Drug Indications were annotated via Broad Institute Connectivity MAP: Drug Re-Purposing hub.

## Discussion

Here we report the largest meta-analysis of GCA using MTAG that adjusted for overlapping samples in the two largest GWAS of cognitive function yet. At an estimated sample size of approximately 373,617individuals, we identified 241 significant genetic loci, of which 39 are novel to the input GWASs, and 29 of these were not reported to be associated with GCA previously. The results are not surprising, in that the original sample overlap between the two reported GWAS were sufficiently large (89%). Consistent with earlier reports of GCA, gene property analysis revealed significant tissue expression overrepresented in GTEX v7/Brainspan brain related tissue compared with expression in other types of tissue. It is notable that some of these genes appear to be significantly expressed during the prenatal state, indicating a potential neurodevelopmental impact of genes that are associated with GCA. In the current study, we focused on identifying genes associated with GCA that could be “actionable” in terms of identifying pharmacological agents that could be re-purposed for nootropic utilization based on GWAS. We have earlier used a similar approach using MAGMA pathway analysis against drug-based pathway annotations on a smaller GWAS of GCA^10^, where we reported several T and L-type calcium channels as potential targets for nootropic agents. Here, we were able to leverage several novel developments: an expanded genome-wide analysis of GCA; newly available brain eQTL· data and complementary transcriptomic methodologies, enabling estimation of directionality (i.e., up-vs. down-regulation of expression) of gene effects on cognition. Notably, our study is the first cognitive GWAS to employ HEIDI, an approach that allows pleiotropy (either vertical or horizontal) to be differentiated from linkage (A single variant is associated with the trait and with gene expression because it is linked by LD to a second variant. However, whilst the first variant is causally linked with the trait, the second variant is causally linked with gene expression). HEIDI tests against the null hypothesis that a single causal variant affects both gene expression and trait variation, and so HEIDI-significant genes are less likely to be causal and require closer inspection and further biological experiments to unravel any true functional effects of the genes. Therefore, we have filtered gene results based on a nominal threshold of *P_HEIDI_* < 0.01. Additionally, several novel classes of gene sets, such as cell binding, cell metabolism, and cell structure not previously reported as associated with GCA, created an additional pool of genes available for further “druggability” investigation.

The most crucial stage of the current report involved the identification of genes that are potential drug targets. Using filtering methods that were detailed earlier, the 76 potentially “druggable” genes were selected for further annotation. Of these, 16 genes were identified as “most likely druggable” based their predicted function from eQTL results and the CMAP Drug Re-purposing database^61^. These selected genes could be further classified into broad gene classes i) Serotonergic genes ii) Carbonic Anhydrase iii) Phosphodiesterase iv) Ion channel v) Glutamatergic/GABA-ergic and vi) Others (See Table 4). Here, we provide further review of the genes/gene classes associated with GCA that could be targeted to improve nootropic function.

### Serotonergic Genes

The most novel and intriguing finding of the present study is the identification of several serotonergic genes as relevant to cognitive function. These genes were not identified under genomewide significant peaks, but rather emerged using our gene-set annotation strategy; therefore, some caution in interpretation should be exercised. Nevertheless, serotonergic mechanisms in cognition have support from several prior lines of research. For example, reduced serotonin may be linked to cognitive disturbances and certain conditions such as Alzheimer’s disease and mood disorder, and stimulating serotonin activity in depression may be beneficial to cognition independent of general relief of depressive symptoms^66–68^. However, antidepressants typically inhibit the serotonin transporter, while the present results suggest that enhancing its function may have pro-cognitive effects. Perhaps more readily interpretable, results of the present study show generally that upregulation of HTR1D and downregulation of HTR5A are associated with enhanced cognitive function. One popular antidepressant, Vortioxetine, is a 5-HT1D agonist and demonstrates some evidence of pro-cognitive efficacy^69^. The triptans, a class of 5-HT1D agonists used for the treatment of migraine, have also demonstrated initial efficacy in rescuing migraine-induced cognitive deficits^70^. At the same time, antagonizing the 5-HT_5A_ receptor has showed cognitive enhancement in a ketamine-based rat model of cognitive dysfunction and negative symptoms of schizophrenia^71^ While ergot-derived migraine treatments with action at 5-HT5A have not shown evidence of cognitive benefit^72^, these agents tend to have complex actions at multiple serotonin (and other neurotransmitter) receptors.

### Carbonic Anhydrase Genes

The current study is the first to report evidence that carbonic anhydrase genes may be implicated in cognitive function. Carbonic anhydrase activity within the hippocampal neurons modulates GABA-ergic functions, altering sensitivity of the gating function for signal transfer through the hippocampal network^73^. Modifying the function of carbonic anhydrase in animal models improved learning abilities, and possible perception, processing and storing of temporally associated signals^74^. Some early reports have also speculated the role of zinc homeostasis being related to cognitive impairment in Alzheimer’s disease^75–77^. While results of the present study suggest that inhibition of carbonic anhydrase activity may enhance cognition, pharmacologic evidence to date has supported the opposite conclusion. Specifically, carbonic anhydrase activation has shown enhancement in synaptic efficacy, spatial learning, memory, as well as object recognition in rodents^74,78^; in humans, topiramate, a carbonic anhydrase inhibitor has been associated with cognitive deterioration^79^. One mechanism by which carbonic anhydrase inhibition might improve cognitive function is in the context of amyloid pathology, which may be dependent on carbonic anhydrase activity.

### Phosphodiesterase Genes

Phosphodieterases (PDEs) catalyze the only known reaction terminating cyclic nucleotide signals, as such, they are crucial regulators of physiological and pathophysiological mechanisms that underlie these processes. Here, we report that a class of PDE-4s, *PDE4D* and *PDE4C* demonstrate association with cognitive function. PDE4s are expressed in the cerebral cortex, hippocampus, hypothalamus, striatum, dopaminergic neurons within the substantia nigra, and astrocytes^80,81^. Inhibiting PDE4 increases the phosphorylation of CREB and hippocampal neurogenesis propagating antidepressant mimicking and memory-enhancing properties^82,83^.Previously reported evidence implicated PDE4s, in particular, *PDE4D,* in possessing pro-cognitive and neuro-protective properties after the infusion of Rolipram^84,85^. The development of therapeutic indications for Alzheimer’s Disease, Huntington’s disease, schizophrenia, depression and cognitive enhancement continues to be the subject of ongoing research^86,87^. Extensive discussion of *PDE4D,* previously discovered by large scale educational attainment GWAS, is reported elsewhere^88^. In the present study, improved eQTL mapping supports the nootropic function of inhibiting *PDE4D.* The role of *PDE4C* appear to be less understood. Earlier reports indicate that though *PDE4C* is expressed in the brain, limited to the cortex, thalamic nuclei and cerebellum^81^. No evidence currently exists to show if activating *PDE4C* plays a conclusive role in rescuing cognitive deficits^89^.

### Glutamatergic Genes

The role of excitatory glutamatergic and inhibitory GABA-ergic neurons are well researched in their relationship to cognitive function and presence in the brain^90,91^. Glutamate mediates fast synaptic transmission and plays a key role in long term potentiation^92^, synaptic plasticity, learning and memory, and other cognitive functions^93^. Extended glutamate stimulation can be damaging to neurons and give rise to excitotoxicity, regarded as a precursor mechanism to several neurodegenerative disorders^94,95^. Indirect modulation of the glutamatergic system via positive allosteric modulators of AMPAR have shown nootropic properties in laboratory animals and human patients^96–100^. Direct modulation of glutamatergic pathway via antagonists, co-agonizing the glycine site, potentiatiating the activity of agonists via polyamines, neurosteroids, and histamines for purpose of cognitive enhancement has also been explored^101^.

Here, we identified AMPA4 agonists as potential cognitive enhancement agents. Within the CMAP Drug Re-purposing database^61^, we identified Piracetam, a known nootropic as an acetylcholine agonist that appear to have shown evidence for improving cognitive function^102^ via complex glutamatergic and calcium signalling pathways^103^. A counterintuitive result was that down regulation of eQTL for *GRIN2A* was related to higher cognitive function. However, it appears that there has been discussion of how low dose antagonism of glutamatergic receptors (N-methyl-D-aspartate: NMDA-R) might increase excitatory effects of glutamate neurons^104^. Supporting evidence for the precognitive effects of NMDAR antagonists like memantine has also been reported in animal models and humans^105^. Based on existing evidence, we also show that the *GRIN2A* gene might also be indirectly targeted by norepinephrine transporter inhibitors, serotonin-norepinephrine reuptake inhibitors, and calcium channel blockers (Table 4).

### Voltage-gated Ion Channel Genes

Voltage-gated ion channels have originally been studied with respect to etiologies of excitability disorders of the heart and muscles. Nevertheless, there is currently emerging evidence for the role of calcium, sodium and potassium channels in the etiopathologies of neuropsychiatric disorders^106^. Ostensibly, these neuropsychiatric disorders and accompanying cognitive function deficits could be rescued by therapeutics aimed at targeting the underlying putative channelopathies^107^. Voltage-gated calcium channels increase periplasmic calcium concentrations, which triggers a downstream cascade of proteins involving ion channel function, vesicle docking and small molecule transport^108^. Calcium trafficking and signalling play a crucial role in cognitive function^109–116^. There is also evidence to suggest that voltage gated calcium channels are necessary for the function of dopaminergic neurons on mesolimbic and mesocortical regions^117,118^. Prior reports have suggested that blocking L-type calcium channels could be a viable strategy for Alzheimer’s disease but noted the paradoxical effect that these channels also promotes synaptic plasticity and spatial memory^119^.

Here, we identify upregulation of *CACNA2D2* and *CACNG3* genes associated with cognitive function. Though calcium channel genes have been identified previously in both cognitive function and neuropsychiatric disease GWASs, work in identifying reliable compounds for calcium activation is relatively nascent. Existing drugs targeting calcium channel receptors are mainly antagonists There is evidence to suggest that indirect activation of calcium channel genes via activating sarco-/ER Ca2+ ATPase 2 (SERCA) appear to be neuroprotective and enhance cognition and memory in Alzheimer’s mouse model^120^. SERCA resides in the endoplasmic reticulum and its dysregulation is thought to affect cognitive function in Darier’s disease, schizophrenia, Alzheimer’s disease, and cerebral ischemia^121^. In the current report, adrenergic receptor agonists could also potentially play a role in activating calcium channel genes highlighted.

Additional to calcium channels, current results also point to the potential role of the chloride voltage channel gene *CLCN2* as a potential gene target for cognitive enhancement. *CLCN2* plays a crucial role in background conductance, removing excess Cl-ions within pyramidal cells in the hippocampus, and regulates excitability in GABAergic interneurons^122^. Loss-of-function mutations in CLCN2 are associated with leukoencephalopathy^123^, and, controversially, with epilepsy^124, 125^; therefore, it is plausible that activation of *CLCN2* might serve to enhance cognitive function. By contrast, gain of function mutations^126^ are associated with primary aldosteronism and subsequent hypertension, without cognitive impairment. The only drug with such a function identified by CMAP search was lubiprostone^127^, which is utilized for constipation and has unknown activity in the CNS.

### Other Genes

Several genes do not fall into clear categories but nonetheless are crucial in the context of cognitive function, I.e., *DPP4, THRB, PSMA5, DHODH2.* While little is known about potential cognitive functions of *DHODH* or *PSMA5,* we examine *DPP4* and *THRB* below.

### Dipeptidyl Peptidase 4

Dipeptidyl peptidase IV (DPP-IV) is a serine protease is known to inactivate glucagon-like peptide-1 (GLP-1), pituitary adenylate cyclase-activating polypeptide (PACAP) and glucose-dependent insulinotropic peptide (GIP), which gives rise to pancreatic insulin secretion. Inhibition of DPP-IV enzyme activity via the gliptin class of medications has thus been widely utilized as a treatment option for diabetes^128^. However, aside from glucose control, animal studies have shown pro-neurogenic^129^, anti-inflammatory^130^ and neuroplasticity^131^ properties. DPP-4 inhibitors appear to improve glucose control and protect against worsening in cognitive functioning in older patients with type 2 diabetes^131^, and in some cases improve cognitive function^132^. Benefits of DPP-4 inhibition in the post-stroke recovery phase and long-term clinical outcome had also been extensively discussed^133^. Reports have also shown that linagliptin possess neuroprotective properties attributed to elevated levels of incretins in the brain^134^, while sitagliptin appear to regulate synaptic plasticity in AD mice via activating GLP-1 and BDNF-Trkb signaling^135^. Data from the current report suggest that downregulation of *DPP4* in is associated with better cognitive function, and therefore *DPP4* inhibitors have been identified as potential drug repurposing candidates for pro-cognitive investigation.

### Thyroid Hormone Receptor Beta

Thyroid hormones (TH) has a vital function in neurodevelopment and its receptors known to regulate neurogenesis in the hippocampus, hypothalamus and subventricular zone^136–138^. In adults, hypothyroidism is related to depressive-like symptomatology, dementia, memory impairment, and psychomotor deficits^139^. These syndromes are thought to be mediated through serotonergic^140^ and/or catecholaminergic^141^ pathways. Treatment of hypothyroidism improved cognitive performance in a mouse model of Alzheimer’s disease^142^ and patients^143^. Evidence for thyroid hormones implicating learning and memory through synaptic plasticity, neuronal cell differentiation and maturation had also been presented^144^. These evidences converge with the data presented in the current study showing that activating the thyroid hormone beta receptor would potentially yield nootropic effects.

The results here have generated leads for further investigation into potential drug functions and how they might provide nootropic function. There are also limitations to the evidence that we report. First, though the evidence reported comprises the largest and most well-powered MTAG analysis of cognitive function, there continues to be potential to expand sample size to increase power. The modest increase in novel loci reported in the current study could be accounted for by substantial sample overlap in the earlier GWAS reports. Second, identifying eQTL for a particular phenotype is challenging–as with most summary statistics approaches, it is not always possible to directly ascertain that eQTL is necessarily leading to variation in the phenotype^145^. As case in point, S-TissueXcan was used as one of the indicators of gene expression direction. Results of S-TissueXcan are powerful in that they index overall potential of expression of the gene investigated but remains a noisy indicator for expression direction. Nevertheless, due to the modest sample sizes available in the annotation databases, our strategy is reasonable at this stage of advancement of biology. Direct experimentation is required to rule out potential extraneous factors that might be pleiotropic to both phenotypic variation and eQTL effects. Third, the issue of LD within GWAS loci has been remained complex^146^, since there are often many genes that reside in some of the genomic regions.

Here, we attempted to identify functionally relevant genes by examining the convergence across a range of complementary methodologies in order to overcome some of the limitations noted above. In addition, we used the HEIDI test to explicitly exclude genes marked by linkage that might be inaccurately labelled as “causal.” Nevertheless, the challenge of regions of large LD and genes should be addressed in future studies, perhaps incorporating recently developed methods for examining three-dimensional properties of the genome^147^. Though we have focused the discussion of results explicitly on identifying potential targets for nootropic purposes, the converse could also be relevant–where there might be commonly administered drugs that appear to result in cognitive deficits e.g. topiramate^148^, gabapentin^149^, and vinorelbine^150^.

We also observed several counter-intuitive findings with respect to directionality of effects; for example, with respect to carbonic anhydrase inhibition. It is plausible that many molecular functions in the brain observe either a U-shape or inverted U-shape curve, such that effects of up- or downregulation are not strictly linear. Moreover, the results reported here are with reference to MTAG conducted in the general population and might appear to be counterintuitive if interpreted with respect to a disease population. For instance, calcium channel blockers might rescue cognitive impairments in schizophrenia, but blocking calcium channel function in the general population could be detrimental to synaptic function. Or enhancing prothrombin in the general population might offer nootropic effects through microtubule function but would potentially increase neurofibrillary tangles in Alzheimer’s disease. At the same time, our GWAS cohorts included older adults, and some findings may be a function of cryptic pathologic processes occurring in these apparently normal subjects. Further work is necessary to replicate evidence reported here into disease populations, along with more precise data on biological mechanisms underlying cognitive function to ensure that compounds identified as nootropic in a population is indeed applicable in certain other disease contexts.

### Conclusions

We performed the largest MTAG analysis for GCA. Aside from identifying 29 fully novel loci in the current study, the effort has included the most well powered cognitive MTAG analysis for identifying genes that are “druggable” and potential drugs that could be repurposed for nootropic utilization. Gene set analysis identified known neurodevelopmental and synaptic related pathways, but also novel cell structure and binding pathways that appeared to subserve known “druggable” genes. Utilizing multiple chemoinformatic and drug repurposing databases, along with eQTL and GWAS data, we identified Serotoninergic, Carbonic Anhydrases, Voltage-gated Ion channels, Glutamatergic/GABA-ergic, and Phosphodiesterase gene classes contribute to GCA, along with miscellaneous genes such as Diphenyl-Peptidase 4, and others. Our efforts show that within these pathways, specific gene classes coding for cellular components and functions could be targeted for nootropic purposes. Further work is necessary to confirm the role of these genes and receptors, to specify their biological mechanisms influencing cognition, and to consider potential CNS effects (including blood-brain barriers permeability) of the putative nootropic compounds nominated by this approach.

## Supporting information

Supplementary Section

