## Supplementary Section for "Identifying Nootropic Drug Targets via Large-Scale Cognitive GWAS and Transcriptomics"

### **Supplementary Tables**

**Supplementary Table 1 | Candidate SNPs within independent loci**

*Note:* Savage.P : GWAS P-values reported by Savage et al; Savage.Z: GWAS Z-scores from Savage et al; Savage.N: Sample sizes reported by Savage et al; Davies.P : GWAS P-values reported by Davies et al; Davies.Z: GWAS Z-scores from Davies et al; Davies.N: Sample sizes derived from Davies et al; Savage.+/-: effect direction from Z-scores from Savage et al; Davies.+/-: effect direction from Z-scores from Davies et al; mtag.n.lower: lower bound estimated sample sizes by MTAG; mtag.n.upper bound estimated sample sizes by MTAG; mtag.frq: allele frequency based on MTAG; mtag_z: Z-score output from MTAG; mtag_pval: MTAG p-values.

**Supplementary Table 2 | Genomic Loci and Top SNP within loci for MTAG analysis**

*Note:* Savage.P : GWAS P-values reported by Savage et al; Savage.Z: GWAS Z-scores from Savage et al; Davies.P : GWAS P-values reported by Davies et al; Davies.Z: GWAS Z-scores from Davies et al; MTAG.SNP: SNP RSID output from MTAG; MTAG.A1/MTAG.A2: Harmonized allele outputs from MTAG; MTAG.Z: Z-score output from MTAG; MTAG.P: P-value output from MTAG.

**Supplementary Table 3 | Winner's Curse Adjustment via FDR Inverse Quantile Transformation**

*Note*: Z_FIQT_ : FIQT adjusted scores. These are Z-scores downweighted by the FIQT algorithm. We indicate Z scores from GWASs from either Davies or Savage et al; Z_predicted-Davies^_ : These are the adjusted Z scores, assuming no winner's curse in Davies; Z_predicted-Savage^_ : These are the adjusted Z scores, assuming no winner's curse in Savage.

**Supplementary Table 4 | Genetic correlations carried out via LD-hub v1.9.3**

**Supplementary Table 5 | S-TissueXcan Transcriptomic Wide Identified Genes**

*Note:* p.adj: Bonferroni adjusted p-values for S-TissueXcan; p_i_best: smallest p-value for tissue type; t_i_best: tissue that has the smallest p-value; p_i_worst: largest p-value for tissue type; t_i_worst: tissue that has the largest p-value

**Supplementary Table 6 | S-PrediXcan results for brain tissue**

Supplementary Table 6a : S-PrediXcan results : Nucleas Accumbens

Supplementary Table 6b : S-PrediXcan results : Amygdala

Supplementary Table 6c : S-PrediXcan results : Anterior Cingulate Cortex

Supplementary Table 6d : S-PrediXcan results : Cerebellum

Supplementary Table 6e : S-PrediXcan results : Cerebellar Hemisphere

Supplementary Table 6f : S-PrediXcan results : Cortex

Supplementary Table 6g : S-PrediXcan results : Frontal Cortex

Supplementary Table 6h : S-PrediXcan results : Hippocampus

Supplementary Table 6i : S-PrediXcan results : Hypothalamus

Supplementary Table 6j : S-PrediXcan results : Putamen

Supplementary Table 6k : S-PrediXcan results : Consolidated Bonferroni P-value Corrected S-Predixcan Gene List

*Note:* P.adj: Bonferroni adjusted p-values

**Supplementary Table 7 | SMR and HEIDI analysis for SNP-Gene Mendelian** Randomization

*Note:* p_SMR: SMR p-values; p_SMR_multi: multi-variant SMR p-values; p_HEIDI: HEIDI p-values; *_adj: bonferroni adjusted p-values; Bemeta: Brain-eMeta annotations; PsychEncode PEER: PsychENCODE data corrected for Probabilistic Estimation of Expression Residuals; PsychEncode HCP: PsychENCODE data corrected for Hidden Covariates with Prior Knowledge; Gtex: 10 brain regions from GTEx

**Supplementary Table 8 | SMR and HEIDI results FDR corrected Gene Lists**

*Note:* p_msmr: multi-variant SMR p-values; p_heidi: HEIDI p-values; B& H q-value: Benjamini & Hochberg q

**Supplementary Table 9 | FUMA eQTL Mapping of Brain Expressed Genes**

**Supplementary Table 10 | PsychEncode/BrainGVEX Gene Expression QTL**

**Supplementary Table 11 | PsychEncode/BrainGVEX Ribosomal Occupancy QTL**

**Supplementary Table 12 | PsychEncode/BrainGVEX Expression Variation QTL**

**Supplementary Table 13 | PsychEncode/BrainGVEX Splicing QTL**

Supplementary Table 13a | PsychEncode/BrainGVEX Splicing QTL Leafcutter Results

Supplementary Table 13b | VEP annotations for Splicing Variants

Supplementary Table 13c | Splice Variant Gene List for Protein Coding Genes represented in MTAG results

*Note:* FDR: FDR p-values; qtl_p: qtl p-values; mtag_freq: allele frequencies based on MTAG output; mtag_beta: MTAG effect size; mtag_se: standard error of MTAG effect size; mtag_pval: MTAG p-values

**Supplementary Table 14 | MAGMA gene based association test results**

*Note:* P.Adj: Bonferroni adjusted p-values

**Supplementary Table 15 | MAGMA pathway analysis with gene results and druggability tier annotations**

*Note:* Gene SET p-values have been Bonferroni adjusted

**Supplementary Table 16 | Drug Annotations from DGIdb, KI and Finan et al., 2017**

**Supplementary Table 17 | Candidate 'Druggable' Genes with gene expression profiles, and drug-disease indications from CMAP/CLUE**

*Note:* Gene identification approaches: These include various strategies to identify genes that are associated with cognitive ability; Drug Targets: These are obtained from filtered drug targets; S-TissueXcan.GTEX7: significant genes identified by S-TissueXcan; Spredixcan.Brain (post-hoc): Nominally associated genes within S-PrediXcan brain tissues; SMR.Brain.Bonferroni: SMR significant genes after Bonferroni correction; SMR.Brain.FDR (post-hoc): SMR significant genes after FDR correction; HEIDI.(p < 0.01): HEIDI genes at P < 0.01 threshold across annotations; MAGMA.Gene.Chisq: Bonferroni corrected significant MAGMA genes from gene association test; MAGMA.pathways: nominally significant genes within Bonferroni corrected significant MAGMA pathways; FUMA.eQTL.Brain: Significant genes from eQTL mapping via the FUMA pipeline; rQTL.PsychEnc.mvQTL: significant PsychENCODE ribosomal occupancy qtl genes; eQTL.PsychEnc.mvQTL: significant PsychENCODE eqtl genes; evQTL.PsychEnc.mvQTL: significant PsychENCODE expression variation qtl genes; sQTL.PsychEnc.mvQT: significant PsychENCODE splicing qtl genes; QTL directions: these effect directions based on eQTL results; CMAP/CLUE filters: annotations from CMAP-DR database, these include counts of drugs within development phase, (e.g. Launch, Phase1, Phase 2, …), and total drugs for specified indications (e.g. neurology/psychiatry, dermatology, cardiology, …)

**Supplementary Table 18 | CMAP Drug Repurposing Database Annotations for 'High Confidence' Genes**

*Note:* MOA: Mechanism of Action; Drug For: Drug indications; Predicted Function: from eQTL effect directions with general cognitive ability.

### **Extended Methods**

#### **Winner’s Curse Adjustment – FDR Inverse Quantile Transformation**

To check for potential Winner’s Curse in loci originally listed as significant in either Savage et al.,^1^ and Davies et al.,^2^ but no longer significant in the MTAG analysis, we first performed FDR Inverse Quantile Transformation (FIQT) on the Z scores of either of the discovery cohorts. Predicted Z-scores are calculated using the square root of sample size scaling factor as follows:
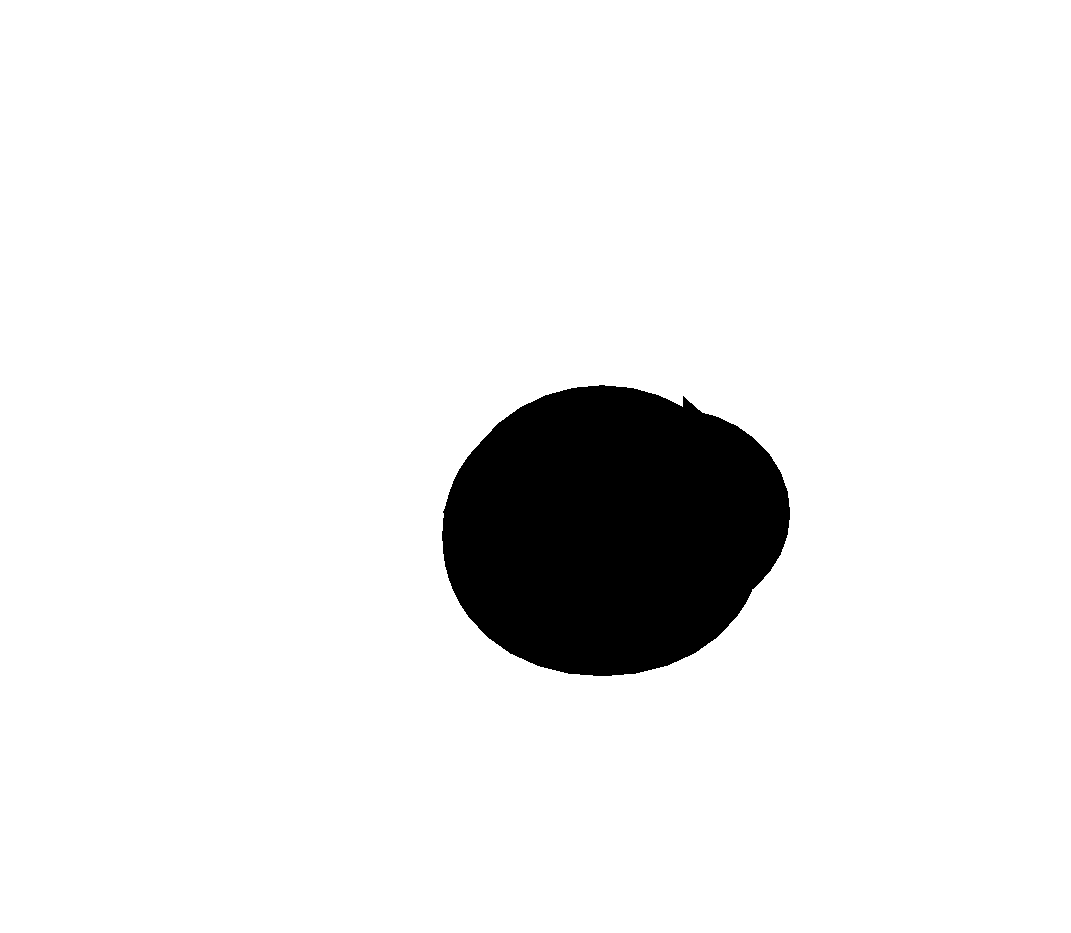

$$Z_{pred}=Z\mu*\surd(\frac{{MTAG}_{N}}{{GWAS}_{N}})$$

Where, $Z_{pred}$ is the predicted Z-scores without Winner’s Curse, $Z\mu$ is the FIQT adjusted Z-scores, and $\surd(\frac{{MTAG}_{N}}{{GWAS}_{N}})$is the scaling factor assuming linear scaling based on sample size increase in fixed-effect meta-analysis as previously described^3^. To test if these results are potentially due to Winner’s Curse, we performed FDR Inverse Quantile Transformation (FIQT) procedures on the input summary statistics. Using simulation methods reports that scales GWAS Z-score effect sizes with sample size, we demonstrated in Supplementary Table 3 that loci that did not meet GWAS significance thresholds were likely related to Winner’s Curse.

### **Acknowledgments**

This work has been supported by grants from the National Institutes of Health (R01 MH117646 to TL; R01 MH079800 and P50 MH080173 to AKM; R01 MH080912 to DCG; K23 MH077807 to KEB; K01 MH085812 to MCK). Data collection for the TOP cohort was supported by the Research Council of Norway, South-East Norway Health Authority, and KG Jebsen Foundation. The NCNG study was supported by Research Council of Norway Grants 154313/V50 and 177458/V50. The NCNG GWAS was financed by grants from the Bergen Research Foundation, the University of Bergen, the Research Council of Norway (FUGE, Psykisk Helse), Helse Vest RHF and Dr Einar Martens Fund. The Helsinki Birth Cohort Study has been supported by grants from the Academy of Finland, the Finnish Diabetes Research Society, Folkhälsan Research Foundation, Novo Nordisk Foundation, Finska Läkaresällskapet, Signe and Ane Gyllenberg Foundation, University of Helsinki, Ministry of Education, Ahokas Foundation, Emil Aaltonen Foundation. We thank the Lothian Birth Cohort participants and research team members. We

thank the staff from the Wellcome Trust Clinical Research Facility at the Western General Hospital

Edinburgh. Phenotype collection in the Lothian Birth Cohort 1921 was supported by the UK

Biotechnology and Biological Sciences Research Council (BBSRC; 15/SAG09977), a Royal Society-Wolfson

Research Merit Award (to Ian Deary), and The Chief Scientist Office of the Scottish Government

(CZH/4/213, CZG/3/2/79, CZB/4/505, ETM/55). Phenotype collection in the Lothian Birth Cohort 1936

was supported by Age UK (The Disconnected Mind project). Genotyping of the cohorts was funded by

the BBSRC (BB/F019394/1, 15/S18386). The work was undertaken by The University of Edinburgh Centre

for Cognitive Ageing and Cognitive Epidemiology (CCACE), part of the cross council Lifelong Health and

Wellbeing Initiative (G0700704/84698, MR/K026992/1), for which funding from the BBSRC and Medical

Research Council (MRC) is gratefully acknowledged. WDH is supported by a grant from Age UK (Disconnected Mind Project). The CAMH work was supported by the CAMH Foundation and the Canadian Institutes of Health Research. The Duke Cognition Cohort (DCC) acknowledges K. Linney, J.M. McEvoy, P. Hunt, V. Dixon, T. Pennuto, K. Cornett, D. Swilling, L. Phillips, M. Silver, J. Covington, N. Walley, J. Dawson, H. Onabanjo, P. Nicoletti, A. Wagoner, J. Elmore, L. Bevan, J. Hunkin and R. Wilson for recruitment and testing of subjects. DCC also acknowledges the Ellison Medical Foundation New Scholar award AG-NS-0441-08 for partial funding of this study as well as the National Institute of Mental Health of the National Institutes of Health under award number K01MH098126. The UCLA Consortium for Neuropsychiatric Phenomics (CNP) study acknowledges the following sources of funding from the NIH: Grants UL1DE019580 and PL1MH083271 (RMB), RL1MH083269 (TDC), RL1DA024853 (EL) and PL1NS062410. The ASPIS study was supported by National Institute of Mental Health research grants R01MH085018 and R01MH092515 to Dr. Dimitrios Avramopoulos. Support for the Duke Neurogenetics Study was provided the National Institutes of Health (R01 DA033369 and R01 AG049789 to ARH) and by a National Science Foundation Graduate Research Fellowship to MAS. Recruitment, genotyping and analysis of the TCD healthy control samples were supported by Science Foundation Ireland (grants 12/IP/1670, 12/IP/1359 and 08/IN.1/B1916).

Data access for several cohorts used in this study was provided by the National Center for Biotechnology Information (NCBI) database of Genotypes and Phenotypes (dbGaP). dbGaP accession numbers for these cohorts were:

Cardiovascular Health Study (CHS): phs000287.v4.p1, phs000377.v5.p1, and phs000226.v3.p1

Framingham Heart Study (FHS): phs000007.v23.p8 and phs000342.v11.p8

Multi-Site Collaborative Study for Genotype-Phenotype Associations in Alzheimer’s Disease (GENADA): phs000219.v1.p1

Long Life Family Study (LLFS): phs000397.v1.p1

Genetics of Late Onset Alzheimer’s Disease Study (LOAD): phs000168.v1.p1

Minnesota Center for Twin and Family Research (MCTFR): phs000620.v1.p1

Philadelphia Neurodevelopmental Cohort (PNC): phs000607.v1.p1

The acknowledgment statements for these cohorts are found below:

**Framingham Heart Study**: The Framingham Heart Study is conducted and supported by the National Heart, Lung, and Blood Institute (NHLBI) in collaboration with Boston University (Contract No. N01-HC-25195 and HHSN268201500001I). This manuscript was not prepared in collaboration with investigators of the Framingham Heart Study and does not necessarily reflect the opinions or views of the Framingham Heart Study, Boston University, or NHLBI. Funding for SHARe Affymetrix genotyping was provided by NHLBI Contract N02-HL-64278. SHARe Illumina genotyping was provided under an agreement between Illumina and Boston University.

**Cardiovascular Health Study**: This research was supported by contracts HHSN268201200036C, HHSN268200800007C, N01-HC-85079, N01-HC-85080, N01-HC-85081, N01-HC-85082, N01-HC-85083, N01-HC-85084, N01-HC-85085, N01-HC-85086, N01-HC-35129, N01 HC-15103, N01 HC-55222, N01-HC-75150, N01-HC-45133, and N01-HC-85239; grant numbers U01 HL080295 and U01 HL130014 from the National Heart, Lung, and Blood Institute, and R01 AG-023629 from the National Institute on Aging, with additional contribution from the National Institute of Neurological Disorders and Stroke. A full list of principal CHS investigators and institutions can be found at https://chs-nhlbi.org/pi. This manuscript was not prepared in collaboration with CHS investigators and does not necessarily reflect the opinions or views of CHS, or the NHLBI. Support for the genotyping through the CARe Study was provided by NHLBI Contract N01-HC-65226. Support for the Cardiovascular Health Study Whole Genome Study was provided by NHLBI grant HL087652. Additional support for infrastructure was provided by HL105756 and additional genotyping among the African-American cohort was supported in part by HL085251, DNA handling and genotyping at Cedars-Sinai Medical Center was supported in part by National Center for Research Resources grant UL1RR033176, now at the National Center for Advancing Translational Technologies CTSI grant UL1TR000124; in addition to the National Institute of Diabetes and Digestive and Kidney Diseases grant DK063491 to the Southern California Diabetes Endocrinology Research Center.

**Multi-Site Collaborative Study for Genotype-Phenotype Associations in Alzheimer’s Disease**: The genotypic and associated phenotypic data used in the study were provided by the GlaxoSmithKline, R&D Limited. Details on data acquisition have been published previously in: Li H, Wetten S, Li L, St Jean PL, Upmanyu R, Surh L, Hosford D, Barnes MR, Briley JD, Borrie M, Coletta N, Delisle R, Dhalla D, Ehm MG, Feldman HH, Fornazzari L, Gauthier S, Goodgame N, Guzman D, Hammond S, Hollingworth P, Hsiung GY, Johnson J, Kelly DD, Keren R, Kertesz A, King KS, Lovestone S, Loy-English I, Matthews PM, Owen MJ, Plumpton M, Pryse-Phillips W, Prinjha RK, Richardson JC, Saunders A, Slater AJ, St George-Hyslop PH, Stinnett SW, Swartz JE, Taylor RL, Wherrett J, Williams J, Yarnall DP, Gibson RA, Irizarry MC, Middleton LT, Roses AD. Candidate single-nucleotide polymorphisms from a genomewide association study of Alzheimer disease. Arch Neurol., Jan;65(1):45-53, 2008 (PMID: 17998437). Filippini N, Rao A, Wetten S, Gibson RA, Borrie M, Guzman D, Kertesz A, Loy-English I,

Williams J, Nichols T, Whitcher B, Matthews PM. Anatomically-distinct genetic associations of

APOE epsilon4 allele load with regional cortical atrophy in Alzheimer’s disease. Neuroimage,

Feb 1;44(3):724-8, 2009. (PMID: 19013250).

**Genetics of Late Onset Alzheimer’s Disease Study**: Funding support for the “Genetic Consortium for Late Onset Alzheimer’s Disease” was provided through the Division of Neuroscience, NIA. The Genetic Consortium for Late Onset Alzheimer’s Disease includes a genome-wide association study funded as part of the Division of Neuroscience, NIA. Assistance with phenotype harmonization and genotype cleaning, as well as with general study coordination, was provided by Genetic Consortium for Late Onset Alzheimer’s Disease. A list of contributing investigators is available at https://www.ncbi.nlm.nih.gov/projects/gap/cgi-bin/study.cgi?study_id=phs000168.v1.p1

**Long Life Family Study: Funding support for the Long Life Family Study** was provided by the Division of Geriatrics and Clinical Gerontology, National Institute on Aging. The Long Life Family Study includes GWAS analyses for factors that contribute to long and healthy life. Assistance with phenotype harmonization and genotype cleaning as well as with general study coordination, was provided by the Division of Geriatrics and Clinical Gerontology, National Institute on Aging. Support for the collection of datasets and samples were provided by Multicenter Cooperative Agreement support by the Division of Geriatrics and Clinical Gerontology, National Institute on Aging (UO1AG023746; UO1023755; UO1023749; UO1023744; UO1023712). Funding support for the genotyping which was performed at the Johns Hopkins University Center for Inherited Disease Research was provided by the National Institute on Aging, National Institutes of Health.

**Minnesota Center for Twin and Family Research**: This project was led by William G. Iacono, PhD. And Matthew K. McGue, PhD (Co-Principal Investigators) at the University of Minnesota, Minneapolis, MN, USA. Co-investigators from the same institution included: Irene J. Elkins, Margaret A. Keyes, Lisa N. Legrand, Stephen M. Malone, William S. Oetting, Michael B. Miller, and Saonli Basu. Funding support for this project was provided through NIDA (U01 DA 024417). Other support for sample ascertainment and data collection came from several grants: R37 DA 05147, R01 AA 09367, R01 AA 11886, R01 DA 13240, R01 MH 66140.

**Philadelphia Neurodevelopmental Cohort**: Support for the collection of the data sets was provided by grant RC2MH089983 awarded to Raquel Gur, MD, and RC2MH089924 awarded to Hakon Hakonarson, MD, PhD. All subjects were recruited through the Center for Applied Genomics at The Children's Hospital in Philadelphia.
